# Allosteric Regulation of Vitamin K2 Biosynthesis in a Human Pathogen

**DOI:** 10.1101/841569

**Authors:** Ghader Bashiri, Laura V. Nigon, Ehab N. M. Jirgis, Ngoc Anh Thu Ho, Tamsyn Stanborough, Stephanie S. Dawes, Edward N. Baker, Esther M. M. Bulloch, Jodie M. Johnston

**Affiliations:** Laboratory of Structural Biology, School of Biological Sciences and Maurice Wilkins Centre for Molecular Biodiscovery, University of Auckland, Private Bag 92019, Auckland, New Zealand; School of Physical and Chemical Sciences, Biomolecular Interaction Centre (BIC), and Maurice Wilkins Centre for Molecular Biodiscovery, University of Canterbury, Christchurch 8041, New Zealand

**Author notes:** Corresponding author: Jodie Johnston, E–mail.

**Keywords:** *Mycobacterium tuberculosis*, Pyruvate oxidase (POX) enzymes, SEPHCHC synthase (MenD), (Thiamine diphosphate) ThDP-dependent enzymes, allostery, feedback inhibition, menaquinone (vitamin K2)

## Abstract

Menaquinone (Vitamin K2) plays a vital role in energy generation and environmental adaptation in many bacteria, including *Mycobacterium tuberculosis* (*Mtb*). Although menaquinone levels are known to be tightly linked to the redox/energy status of the cell, the regulatory mechanisms underpinning this phenomenon are unclear. The first committed step in menaquinone biosynthesis is catalyzed by MenD, a thiamine diphosphate-dependent enzyme comprising three domains. Domains I and III form the MenD active site, but no function has yet been ascribed to domain II. Here we show the last cytosolicmetabolite in the menaquinone biosynthesis pathway (1,4-dihydroxy-2-napthoic acid, DHNA) binds to domain II of *Mtb*-MenD and inhibits enzyme activity. We identified three arginine residues (Arg97, Arg277 and Arg303) that are important for both enzyme activity and the feedback inhibition by DHNA: Arg277 appears to be particularly important for signal propagation from the allosteric site to the active site. This is the first evidence of feedback regulation of the menaquinone biosynthesis pathway in bacteria, unravelling a protein level regulatory mechanism for control of menaquinone levels within the cell.

## Introduction

*Mycobacterium tuberculosis* (*Mtb*), the causative agent of tuberculosis in humans, is able to adopt a persistent phenotype, resulting in long treatment times and a hard-to-eradicate latent infection [1]. To combat this latent state, there has been a growing interest in menaquinone (vitamin K2, MK), a small redox molecule that is essential for energy generation in both actively growing and persistent *Mtb* [2]. MK also plays a role in triggering persistence in *Mtb* through its capacity to signal redox status [3]. Previous studies have shown that inhibiton of MK biosynthesis enzymes can significantly reduce growth of persistent-state like and drug-resistant *Mtb* [4–6]. Therefore, a fundamental understanding of the MK biosynthesis pathway and its regulatory mechanism would provide a deeper insight into the underlying complexity of *Mtb* biology, opening up novel approaches for anti-TB therapeutics.

MK levels are known to be tightly linked to the redox potential in bacteria [2, 3, 7–10]; however, the molecular mechanisms that regulate this phenomenon are unclear. The first committed step in MK biosynthesis in *Mtb* is catalyzed by the thiamine-diphosphate (ThDP)-dependent enzyme MenD (2-succinyl-5-enolpyruvyl-6-hydroxy-3-cyclohexadiene-1-carboxylate synthase, SEPHCHC synthase). Like other members of the ThDP-dependent pyruvate oxidase (POX) family, which are dimers or tetramers (comprising two interfacing dimers), MenD is tetrameric, with each monomer comprising three domains [11, 12]. Domains I and III have known roles in catalytic function, domain I from one monomer in the dimer pairs with domain III of the other monomer (and vice versa) to form two paired active sites per dimer, with residues from both domains contributing to each active site [13–15]. Domain II, however, is much less conserved and does not appear to participate in cofactor or substrate binding [11, 15]. Various roles have been suggested for domain II in other POX-family enzymes; for example, substrate and substrate-like allosteric activators bind in domain II of pyruvate decarboxylase (PDC) and phenylpyruvate decarboxylase [11, 12, 14–18], and nucleotides bind in other enzymes (e.g. FAD in POX and ADP in oxalyl CoA decarboxylase) [11, 19–21]. However, the function of domain II in MenD remains unexplored.

We previously determined a series of crystal structures of *Mtb*-MenD showing each step in the MenD catalytic cycle, as substrates α-ketoglutarate and isochorismate are successively added to the ThDP cofactor before the final product is released [13]. We have now identified a downstream metabolite of the MK biosynthesis pathway (1,4-dihydroxy-2-napthoic acid, DHNA) that binds to domain II of *Mtb*-MenD and inhibits its catalytic activity. Herein we characterize DHNA binding to *Mtb*-MenD at the molecular level, providing evidence for protein-level allosteric regulation and feedback inhibition of the classical MK biosynthesis pathway in *Mtb* and opening up new and unanticipated possibilities for therapeutic intervention in this important pathway.

## Results and Discussion

### A downstream metabolite of the MK biosynthesis pathway binds to domain II of *Mtb*-MenD

The classical MK biosynthesis pathway (Figure 1A) starts with the synthesis of the napthoquinone head-group precursor (DHNA) in the cytosol [22, 23], followed by prenylation and methylation by membrane bound enzymes to produce the lipid soluble MK [24, 25]. In addition to its electron transport role, the prenyl tail of MK can be further modified and these modified quinones have been shown to regulate virulence of the *Mtb* infection [7, 26, 27]. *Mtb*-MenD is situated at a key step in MK biosynthesis, the first committed step (Figure 1A/B) [28, 29] and we hypothesized that it might be involved in controlling flux though this pathway. In particular, we aimed to determine whether *Mtb*-MenD is subject to feedback regulation. We soaked crystals of *Mtb*-MenD with a range of substrates, metabolites, and metabolite-like compounds from the MK biosynthesis pathway to determine whether any of them bound to the enzyme, and found that DHNA, a downstream metabolite of MK biosynthesis, binds to a site in domain II of *Mtb*-MenD (Figure 1C).

**Figure 1.**
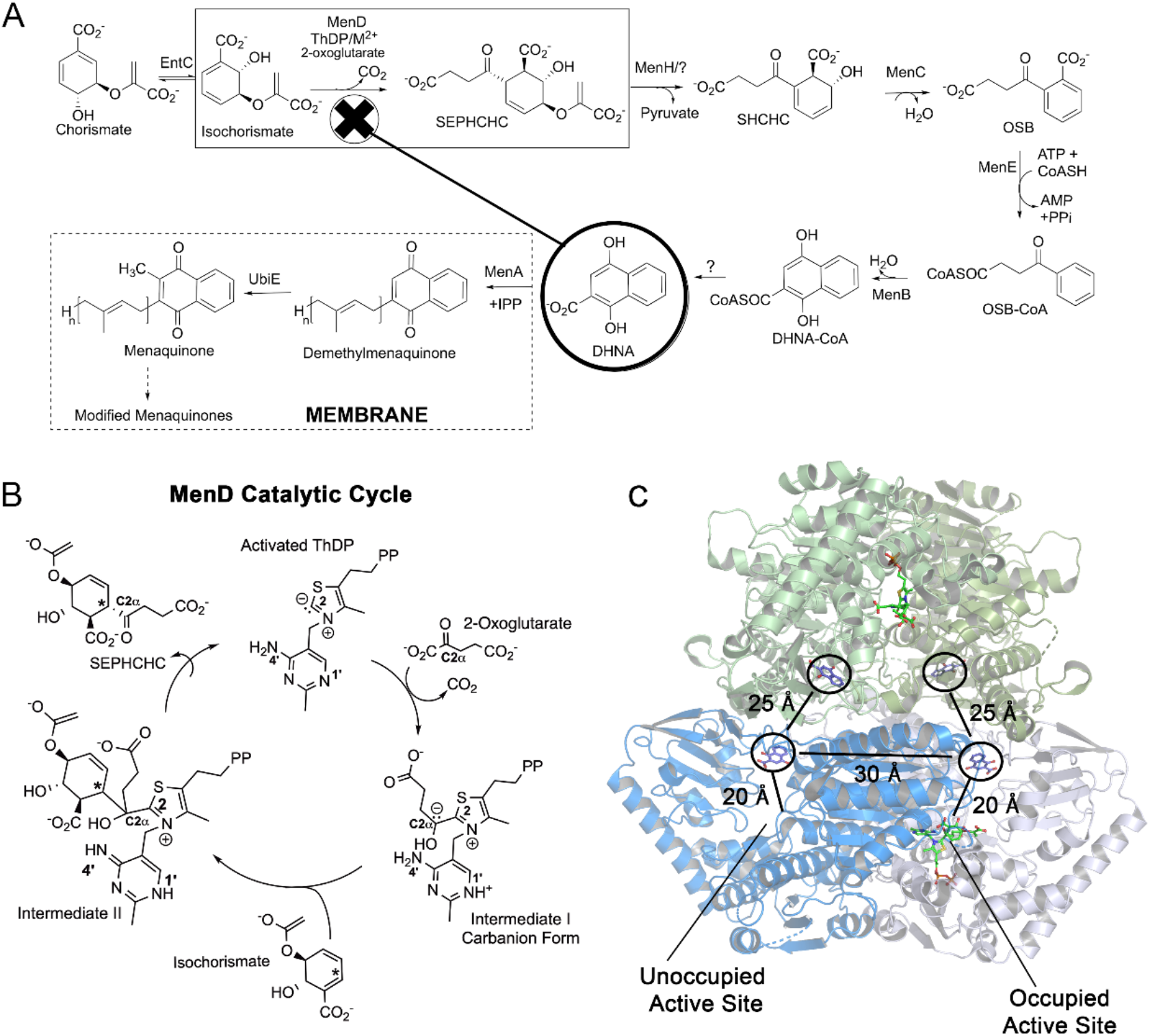
The MK Biosynthesis pathway showing binding of DHNA to *Mtb*-MenD. **A.** Known enzymes in the classical biosynthesis pathway. MenD is the first committed step, with the step before often carried out by an isochorismate synthase enzyme non-specific to the pathway (e.g. EntC in *Mtb*). DHNA is the last metabolite produced by a cytosolic enzyme. **B.** The MenD catalytic cycle showing the two covalent ThDP intermediates. **C.** The *Mtb*-MenD tetramer (composed of two interfacing dimers, one depicted as green/light green cartoons and the other as blue/pale blue cartoons). DHNA is shown as blue sticks and intermediate II as green sticks. The approximate distance of DHNA to the closest allosteric sites (which is across the dimer-dimer interface) and closest active sites (within the same dimer, shown for a single dimer only for clarity) are depicted by lines/distance labels.

To further characterize the interactions between *Mtb*-MenD and DHNA we determined four crystal structures representing different functional states of the enzyme; an apo (cofactor-free) form complexed with DHNA, acofactor (ThDP) bound DHNA complex and two reaction intermediate-bound (intermediate I and II, Figure 1B) DHNA complexes **(Supplementary Table 1)**. DHNA binding to *Mtb*-MenD was consistently observed in all four subunits of each structure, i.e. with and without the ThDP cofactor, and in the reaction intermediate-bound forms. In all cases, DHNA bound between domains I and II of each MenD subunit, in a cleft formed by residues 94-97, 232-235, 276-278 and 299-306 and capped by residues 111-117 from a neighbouring subunit (Figure 2A). This cleft is located ∼ 20 Å away from the closer of the paired active sites in the dimer and ∼ 30 Å from the more distant one (Figure 1C and 2B).

**Figure 2.**
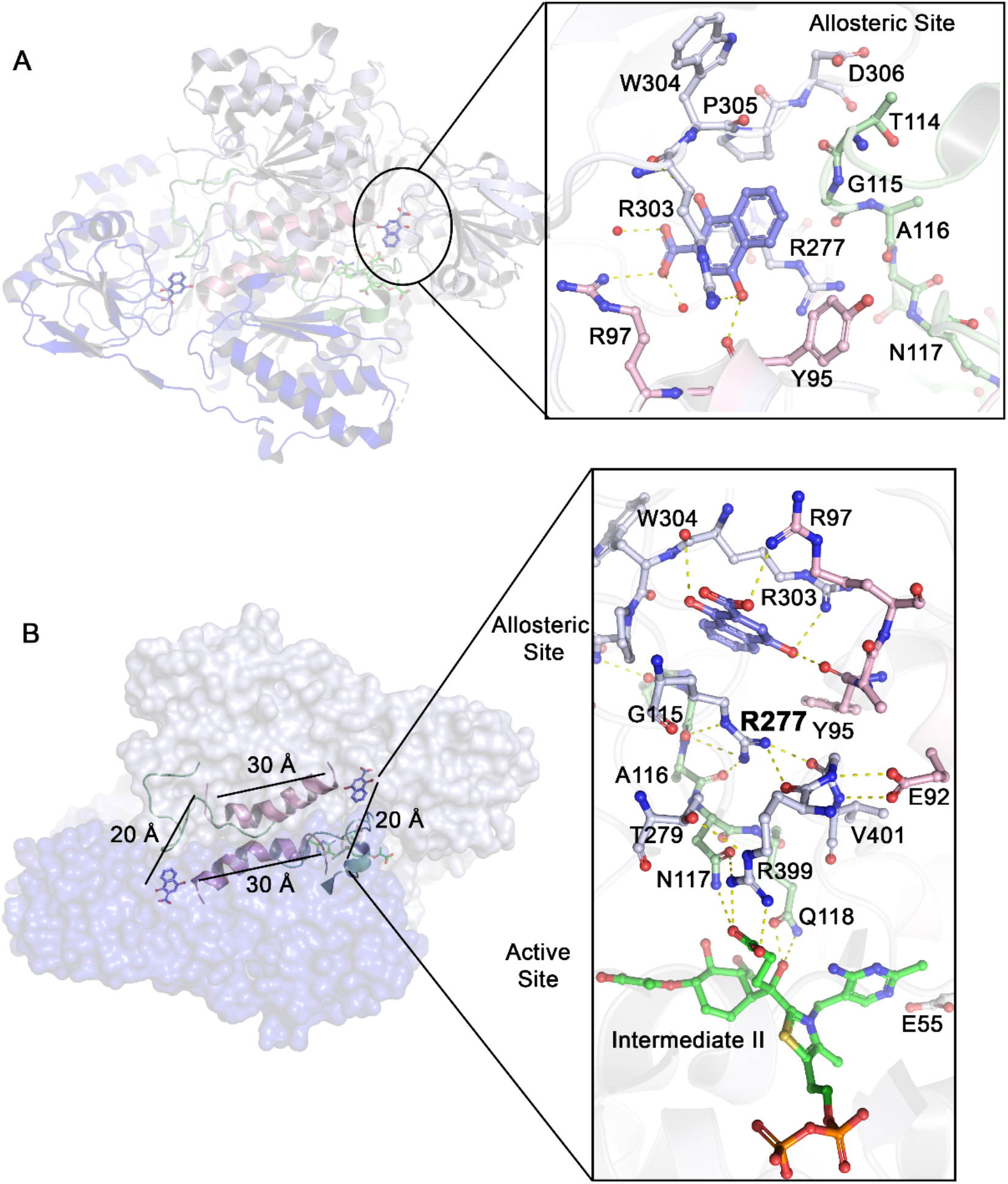
The DHNA binding site is connected to the active site. **A.** The DHNA binding sites from one of the *Mtb*-MenD dimers, with a close up of the DHNA binding site nearest to an occupied active site. The binding site is surrounded by residues from domain II (pale white blue sticks) and domain I (residues 95-97 shown as pink sticks) regions from one monomer in the dimer, and the site is capped by residues 111-116 of the domain I active site loop 105-125 of the neighboring monomer in the dimer (pale green sticks). **B.** Close up of the same dimer showing the network between the allosteric site and its closest active site. The hydrogen bonding network between the two sites from the DHNA binding region through to key intermediate II binding residues is shown. Arg277, with its interactions with two important active site regions (105-125 loop and 399-401) is highlighted in bold.

The binding site for DHNA (Figure 2A) is essentially the same in all four structures. The DHNA molecule occupies an ‘arginine cage’ formed by three arginine residues, Arg97, Arg277 and Arg303, arranged such that the side chains of Arg277 and Arg303 pack on either side of the planar dihydroxynaphthoic acid ring and the side chain of Arg97 hydrogen bonds to the DHNA carboxylate. Additional hydrogen bonding interactions are made by the DHNA hydroxyl groups with the carbonyl oxygens of Tyr95 and Arg303 and the Arg303 guanidinium group. The binding cleft is closed off by residues 114-120 from a flexible active site loop belonging to the other subunit of the MenD dimer; Gly115 makes van der Waals contacts with the DHNA ring.

Whether there is any connectivity between the four allosteric DHNA binding sites in the *Mtb*-MenD tetramer (as there is between the active sites [13]) is unclear. However, the domain II residues 299-306 at the DHNA binding site are connected by a hydrogen bonding network involving residues Arg97, Ala170, Arg159 and Arg168 to the same region in a neighboring *Mtb*-MenD subunit ∼25 Å away. This suggests that binding events on one subunit could be transmitted to the others through such a network.

The location of the DHNA binding site is similar to that of allosteric activator pockets present in another member of the POX superfamily, pyruvate decarboxylase [30]; therefore, we carried out assays to determine whether DHNA acts in a similar manner, as an allosteric regulator of MK biosynthesis.

### DHNA is a potent inhibitor of *Mtb*-MenD SEPHCHC synthase activity

To investigate whether DHNA plays a regulatory role in MK biosynthesis, we studied its effect on *Mtb*-MenD activity. An NMR-based activity assay, similar to that previously reported [13], was first used and showed that the activity of *Mtb*-MenD at a concentration of 5 µM was reduced in the presence of 20 µM DHNA to only 24% of its non-inhibited activity (Figure 3A). Further increases in DHNA concentration resulted in only small increases in inhibition (data not shown), consistent with saturation of the enzyme at low micromolar concentrations.

**Figure 3.**
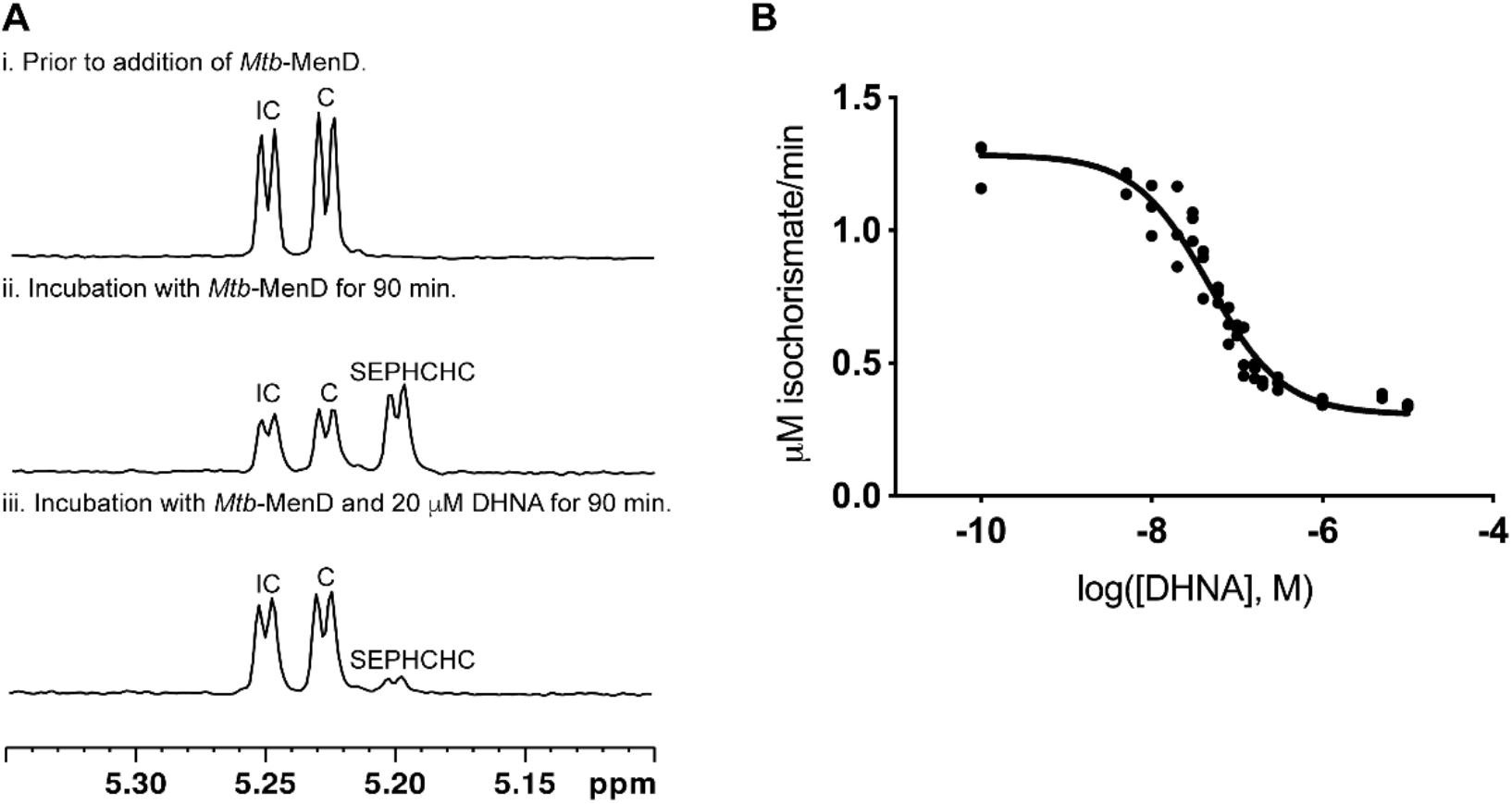
Inhibition of *Mtb*-MenD by DHNA. **A.** Activity assays monitored by ^1^H NMR spectroscopy showing evidence for inhibition of *Mtb-*MenD in the presence of DHNA. Initial reactions solutions, (i) containing an equilibrium mixture of isochorismate (IC, δ 5.25 ppm) and chorismate (C, δ 5.23 ppm) were generated by incubating 2 mM chorismate with 25 µM *E. coli*-MenF for 20 min at 25 °C. The reactions also contained 1 mM α-ketoglutarate and 200 µM ThDP. Subsequently 5 µM *Mtb-*MenD was added and incubation was carried out for a further 90 min at 25 °C either in the absence (ii) or presence (iii) of 20 µM DHNA. In the presence of DHNA the rate of production of SEPHCHC (δ 5.20 ppm) was 24 ± 1% of that in its absence. The doublet peaks assigned to isochorismate, chorismate and SEPHCHC correspond to the equivalent methylene hydrogen (H8b) on the enolpyruvyl group of each compound. Full ^1^H NMR assignment of SEPHCHC was reported previously [13]. **B.** IC_50_ for DHNA against WT *Mtb*-MenD measured using a UV spectrophotometry based-assay for isochorismate consumption. Initial inhibition assay solutions contained 0.6 µM *Mtb*-MenD, 300 µM ThDP and various concentrations of DHNA (0 to 10 µM) and were pre-incubated for 30 min at 25°C. Then 2 µM isochorismate was added and the reaction was initiated by the addition of 300 µM of α-ketoglutarate. Initial rates were measured and fit to the four-parameter logistic Hill equation (solid line).

A UV spectrophotometry-based assay, in which the consumption of isochorismate is monitored at 278 nm, was then used to examine the inhibition over a lower DHNA concentration range (0.1 nM to 10 µM). We found that DHNA inhibited *Mtb*-MenD with an IC_50_ of 53 nM under the conditions of this assay (Figure 3B, Table 1). In combination with our structural complexes, these assays establish DHNA as a potent allosteric inhibitor of *Mtb*-MenD.

**Table 1.** Inhibition of MenD constructs by DHNA.

| MenD variant | Activity<br>relative to<br>wild type (%) | DHNA<br>IC<sub>50</sub> (nM) | 95% confidence<br>interval for<br>DHNA IC<sub>50</sub><br>(nM) | Hill slope |
| --- | --- | --- | --- | --- |
| Wild type | 100 | 53 | 43 - 63 | -1.5 ± 0.2 |
| R97A | 50 | 1030 | 580 - 2700 | -0.8 ± 0.2 |
| R277A | 18 | n.d. | n.d. | n.d. |
| R303A | 56 | 320 | 210 - 490 | -1.0 ± 0.2 |

### Connectivity between the allosteric and active sites of *Mtb*-MenD

How might binding at a site that is remote from the active site impact on catalysis? *Mtb*-MenD is a complex enzyme, characterized by significant conformational changes and disorder-order transitions that take place during the catalytic cycle [13]. In its apo (cofactor-free) state there are substantial regions of disorder. Cofactor-bound structures, including the two covalent intermediates, are more ordered and are also asymmetric, with only two of the four active sites occupied per tetramer [13]. The most striking conformational change associated with DHNA binding involves a flexible active site loop, residues 105–125, that carries the substrate binding residues Arg107, Asn117 and Gln118. This loop is disordered in the apo state, but becomes fully ordered when its associated active site is occupied (Figure 4A). Also reorganized are residues 79-82 at the N-terminus of an α-helix that helps form the binding pocket for the 4’-aminopyrimidine (AP) ring of the ThDP cofactor. These changes are required to generate the catalytically-competent state of the enzyme, and presumably also to enable product release.

**Figure 4.**
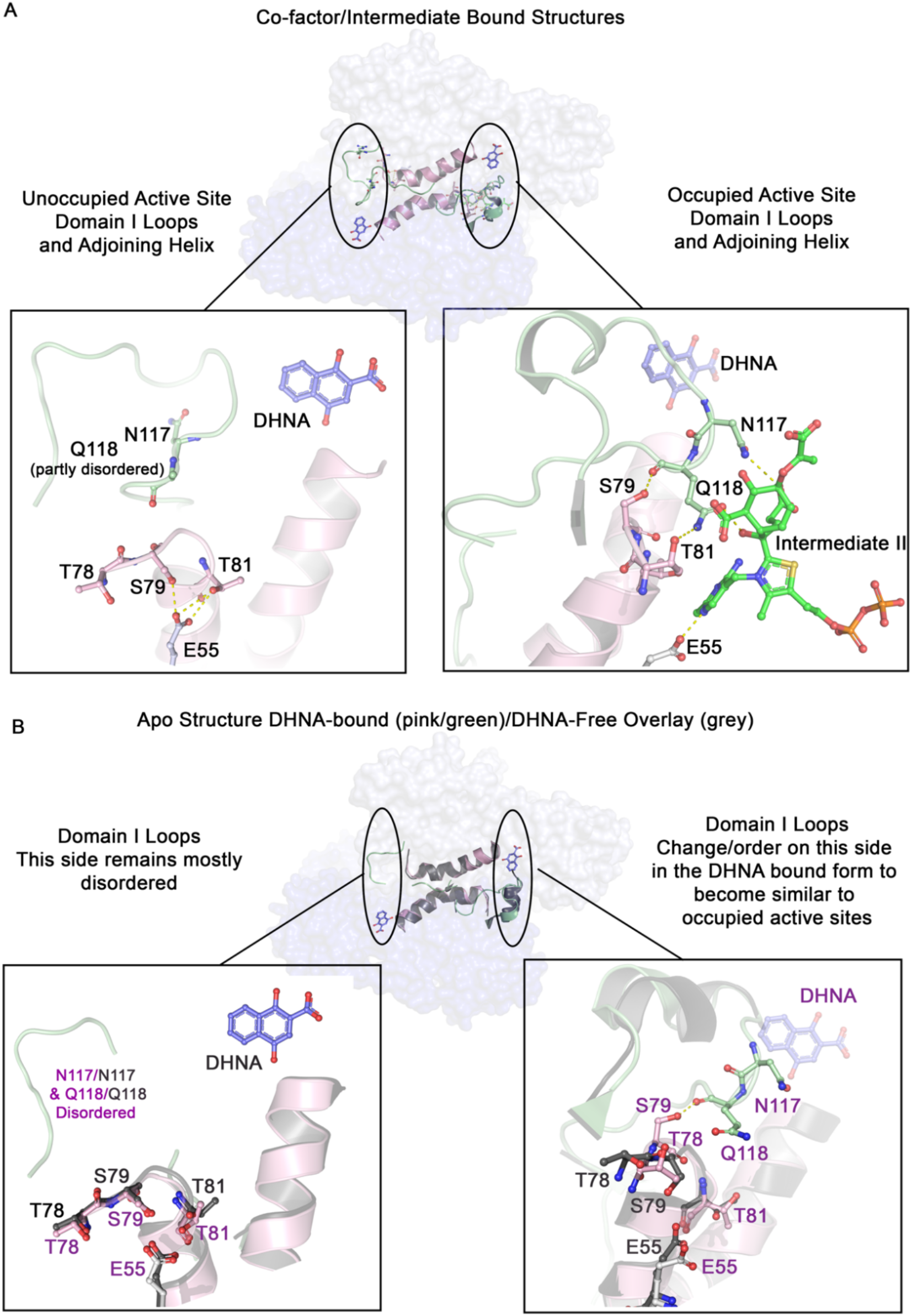
Ordering of two domain I loops. **A.** Typical conformation for the occupied and unoccupied active sites of a dimer in apo-factor/intermediate bound *Mtb*-MenD tetramer. DHNA is shown as blue sticks, intermediate II as green sticks. The domain I 105-125 loop is shown in green with Asn117 and Gln118 depicted as sticks. The 78-97 domain I loop and helix is in pink with Thr78, Ser79 and Thr81 as pink sticks. Catalytic Glu55 is shown as white sticks. In one site Glu55 is bound by Thr81 and Ser79, in the other site these residues move away from Glu55 toward Gln118. **B**. The same view as for A but with overlaid dimers from an APO DHNA-free (grey) and APO DHNA-bound structure (pink/white/green) showing the movements in the 105-125 (green) and 78-82 (pink) loops in the DHNA bound structure.

The DHNA binding site is ∼20 Å from the closest active site, with the two sites separated primarily by two sections of polypeptide, residues 399-402 from domain III and residues 114-120 from the flexible active site loop contributed by the other subunit of the dimer (Figure 2B). Residues 114-120 carry two key active site residues, Gln118, essential for catalysis [13], and Asn117, which binds to the α-ketoglutarate moiety when intermediate I is formed. Similarly, the 399-402 loop provides Arg399, critical for α-ketoglutarate recognition, and Ala402, whose carbonyl oxygen hydrogen bonds to the ThDP AP ring (Figure 2B). DHNA binding to the apo-enzyme causes residues 114-120 to become fully ordered as they complete its binding site, and is associated also with reorganization of residues 78–82, as a new hydrogen bond is made between Ser79 and Gln118 (Figure 4B). The hydrogen bonding environment of the catalytically-essential Glu55 is also changed as a result, to a state part-way between that seen in occupied and unoccupied active sites. This occurs for two of the four active sites creating asymmetry previously only observed in co-factor occupied structures. In many respects the structural changes that occur as DHNA binds to the apo-enzyme thus mirror key changes in domain I that occur when ThDP binds.

From the above it is clear that there is connectivity between the two sites such that the allosteric site is affected by, and can influence, the active site of *Mtb*-MenD. How, then, could DHNA binding cause inhibition of enzyme activity? From this work, the clearest effect of DHNA appears to be to induce asymmetry in apo-MenD and lock in place flexible elements of the active site contributed by domain I into a conformation similar to that seen when ThDP is bound. There are also connections from the allosteric site to domain III components of the active site, (i.e. Arg399) although no domain III ordering/conformational changes are observed, and the flexible domain III loop 471-486 does not adopt its closed form until ThDP is bound. Due to the asymmetry in these domain I and III regions, they are candidates for any order-disorder transitions that might play a role in orchestrating the half-of-sites occupancy. DHNA binding could thus impact any communication between the active sites, as well as many points along the catalytic cycle. The two domain I regions most affected have roles ranging from direct binding to substrates and covalent intermediates to more indirect roles positioning other elements in the active site. As substrate binding, formation of intermediates and product release (both CO_2_ and the final SEPHCHC) all depend on flexibility in the active site (13), DHNA binding, by limiting active site flexibility, has the power to affect catalysis.

A key player in this scenario is likely the invariant Gln118, even conservative mutations of which abolish SEPHCHC synthase activity [31]. In unoccupied active sites, Gln118 is disordered or has high B factors. In occupied sites, however, it is ordered but undergoes sidechain movements that enable specific interactions that are critical to several key steps of the catalytic cycle. These include stabilizing the active tautomer of the AP ring and hydrogen bonding to both the incoming isochorismate substrate and then the resultant intermediate II. It also interacts with the “CO_2_-like” formate ion that likely models the location of the carboxyl group that is removed during formation of intermediate I [13, 32].

### The three arginine cage residues support inhibition by DHNA and dramatically affect enzyme activity

The three arginine residues (Arg97, Arg277, and Arg303) that form a cage around DHNA in its binding site are candidates for signaling between the allosteric and active sites. All three residues interact directly with DHNA and are likely to enhance binding (Figure 2A/B). Arg97 is located at the C-terminus of a long helix that originates in the more distant of the paired active sites, indicating a potential line of communication with that site (Figure 2A/B). Arg277 hydrogen bonds with two regions of the closer of the paired active sites, i.e. with Gly400/Arg399, and with two residues from the 105–125 active site loop (Figure 2B). These regions contain residues (Arg107, Asn117, Gln118, and Arg399) known to be important for *Mtb*-MenD function [28, 32]. Arg303 is located in a region (residues 299-308) that interacts with another part of the 105-125 loop (Figure 2A) and is also involved in a hydrogen bonding network with the closest allosteric site across the tetramer.

To test the importance of these three residues for either MenD activity and/or DHNA inhibition, we carried out alanine mutagenesis experiments. These confirmed that each of the three arginines is crucial for MenD activity (Table 1): the R303A, R97A, and R277A mutant proteins had 56%, 50%, and 18%, of the wild type (WT) enzyme activity, respectively, when measured under the same conditions and in the absence of DHNA. In terms of DHNA inhibition, R97A and R303A showed 19-fold and 6-fold increases in IC_50_, respectively, compared to WT *Mtb*-MenD (Table 1) indicating the importance of these arginines for DHNA binding and feedback inhibition. The R277A variant had such a low catalytic activity that its IC_50_ could not be measured. We conclude that these three residues are important for maintenance of WT MenD activity, probably because they stabilize other elements of the active site (e.g. the 105-125 loop) and that this underpins their roles in signal propagation from the allosteric site to the active site of *Mtb*-MenD.

### How conserved is this allosteric site in other MenD enzymes?

To explore how widely conserved the newly discovered allosteric site is across the bacterial kingdom, we analysed both sequence and structural conservation of this site within MenD orthologues (Figure 5B/5C). Superpositions of the known *E. coli (Ec), Listeria monocytogenes* (*Lm*) and *Bacillus subtilis* (*Bs*) MenD structures [28, 31–34] on to *Mtb*-MenD (overall root-mean-square differences (RMSDs) of 1.8–2.4 Å over 406– 477 aligned Cα atom positions) suggest limited conservation of the DHNA binding site. In *Ec*-MenD and *Lm*-MenD few of the allosteric site residues are conserved and, in *Ec*-MenD, the site is partly filled by the hydrophobic Leu316. In *Bs*-MenD the site retains some key elements including two of the three arginine-cage residues; *Bs*-Arg96 (equivalent to *Mtb-* Arg97) and *Bs*-Arg323, adjacent to the *Bs*-Trp322 (equivalent to *Mtb*-Trp304), which, with rearrangement, could fill an equivalent role to *Mtb*-Arg277 (Figure 5B). A multiple sequence alignment of 35 different bacterial, archeal and plant MenD amino acid sequences using MAFFT [33] (**Supplementary Figure 1**) shows that conservation across domain II is generally low and the three key DHNA binding arginines (Arg97, Arg277, and Arg303) are present only in closely related species such as other *Mycobacteria* (e.g. *Mycobacterium canetti*) and *Rhodococcus* (overall 66.1% sequence identity to *Mtb*-MenD). A further six of the 35 sequences showed conservation for two of the three arginines (Arg97 with either Arg 277 or Arg303) or had Arg97 and an arginine adjacent to the Trp304 eqivalent residue as observed for *Bs*-MenD.

**Figure 5.**
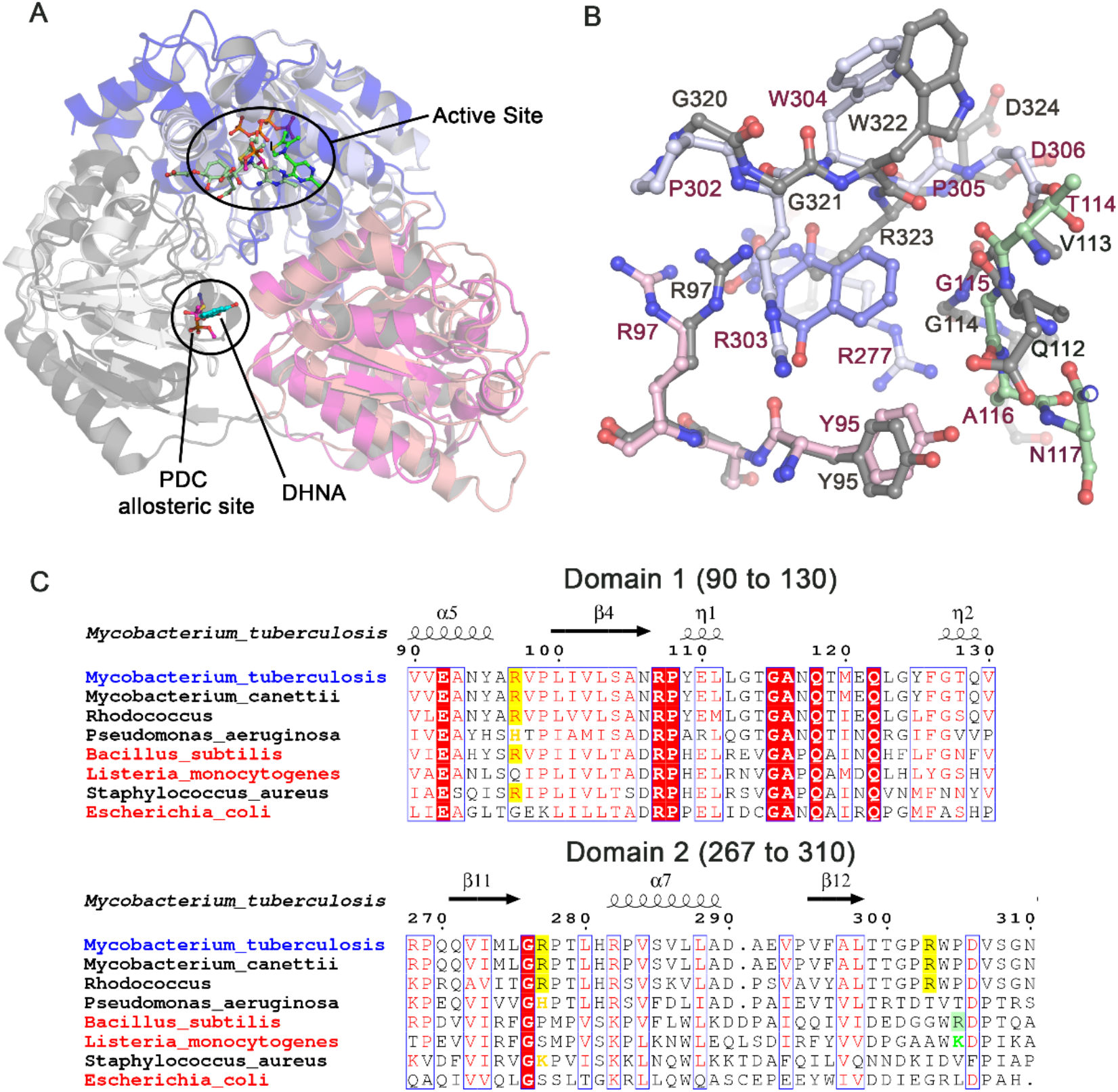
Conservation of the allosteric site. **A.** Location of the DHNA binding site in *Mtb*-MenD compared to the substrate modified allosteric site in pyruvate decarboxylase (PDC, right). Domain I of each structure is shown in pink (*Mtb*-MenD) and magenta (PDC), domain II in white/blue (*Mtb*-MenD) and grey (PDC) and domain III in light blue (*Mtb*-MenD) and deep blue (PDC). ThDP or ThDP intermediates, DHNA and modified Cys-221 are shown as sticks and labelled. **B.** Overlay of the allosteric site of *Mtb*-MenD (pink, light grey and green) with the equivalent regions in *Bs*-MenD (grey). The *Bs*-MenD site shows some conservation, but with no clear Arg303 equivalent. *Bs*-Arg323 could, if the sidechain rearranged take up a similar position to *Mtb*-Arg277. **C.** Representative sequence alignment of domain I and domain II regions of MenD enzymes that contain the Arg allosteric residues (yellow boxes) generated by ESpript[52]. The three sequences with representative structure in the PDB are named in red. The *Bs*-Arg323 position is shown in a green box. Residues similar to Arg in each position (i.e. His or Lys) are coloured in orange or green.

Our studies suggest there is limited strict conservation of key elements of the allosteric site across bacterial MenD enzymes, which could indicate the regulation of the pathway by DHNA is limited to a small sub-set of bacteria. However, considering the partial conservation in some organisms, it is possible that DHNA or related molecules could still bind in this region despite the absence of some key residues found in *Mtb*-MenD. Exactly how widespread allosteric regulation of menaquinone biosynthesis is among bacteria, therefore, remains a question for future work. However the presence of a site with a powerful ability to regulate enzyme activity is of immediate value as a target for species-specific antimicrobials.

### Domain II is adapted for regulatory roles in other ThDP-dependent enzymes

Our results demonstrate that *Mtb*-MenD is an allosterically-regulated ThDP-dependent enzyme, inhibited by direct binding of a downstream metabolite (DHNA) from the biosynthetic pathway in which it participates. While feedback inhibition of this type has not been demonstrated for ThDP-dependent enzymes before, there is precedence for various mechanisms of allosteric regulation in the wider superfamily. Some enzymes, such as acetohydroxyacid synthase, have an entirely distinct negative regulatory subunit [34]. Others, such as 2-ketoglutarate dehydrogenase (KGD) and 2-hydroxy-3-oxoadipate synthase, are allosterically activated by binding of a small molecule (acetyl-CoA) to a shallow pocket on the enzyme surface and allosterically inhibited by binding of a regulatory protein (GarA) [35–37]. Similarly, pyruvate decarboxylase [30], phenylpyruvate decarboxylase [17, 18], and oxalyl-CoA decarboxylase [21] have all been shown to be positively regulated by the direct binding of small molecules to domain II.

Structural comparisons of DHNA-bound *Mtb*-MenD reveal striking parallels with pyruvate decarboxylase, which is allosterically activated by its pyruvate substrate [30]. The pyruvate decarboxylase allosteric site has previously been described as a “switch point” in domain II [30] and has a very similar structural context to the DHNA-binding allosteric site we observe in *Mtb*-MenD (Figure 5A).The location of the allosteric sites map to each other; DHNA overlays with the susbtrate bound in the pyruvate decaboxylase allosteric site. The two activation loops in pyruvate decarboxylase (residues 288–304 and 104–113) that undergo conformational change upon allosteric regulator binding are equivalent to regions in *Mtb*-MenD that are affected by DHNA binding (i.e., residues 277–312 and residues 105–116). Moreover, the substrate binding residues Asn117 and Gln118 in *Mtb*-MenD, which are positioned in the active site when the 105–125 loop is ordered, align with two active site histidines (His114 and His115) in pyruvate decarboxylase [30] that also rearrange when the allosteric site is occupied.

Despite these similarities, key differences exist. The allosteric effect in pyruvate decarboxylase is activation not inhibition, and, in keeping with this, the effector of *Mtb-*MenD is not the substrate but a downstream metabolite. In addition, while the asymmetric dimers of pyruvate decarboxylase undergo large quaternary conformational changes [38], no significant alterations in quaternary structure have been observed for any MenD structures reported to date. The enzymes also diverge in function, catalyzing reactions with different sized substrates and requiring different dynamics during their catalytic cycles [18, 30]. Nonetheless, these parallels do point to the wider use of domain II as a regulatory domain for allosteric regulation of ThDP-dependent enzymes, a phenomenon now also seen in MenD.

### The biological significance of a regulatory role for DHNA

The ability of DHNA to act as a regulatory signal in the pathogen *M. tuberculosis* is in line with both the biological significance of DHNA and the importance of regulating menaquinone levels within the bacteria. As the last non-prenylated soluble metabolite in the MK biosynthetic pathway, DHNA sits at the point where the pathway moves from an aqueous cytosolic location to a lipophilic membrane-immersed one [25] and has the potential to provide feedback on the catalytic status of MenA (and perhaps the downstream MK pool). DHNA is also the first metabolite in the pathway with a complete redox-capable napthoquinone ring [39], and has the capacity in its own right to catalyze redox reactions [40]. It may thus act as a signal of redox status, with excessive levels exerting toxicity if the redox balance within the cell is disrupted. DHNA has also been shown to act as a virulence factor in the intracellular pathogen *Listeria monocytogenes*, where it promotes cytosolic survival, and may be a sensor of cytosolic stress [41]. In plants, phylloquinone biosynthesis enzymes share homology to those of classical bacterial MK biosynthetic enzymes and 1,4 naphthoquinones derived from DHNA act in roles mediating plant–plant, plant–insect and plant–microbe interactions [42].

## Conclusions

Our study reveals the downstream metabolite DHNA as a negative allosteric regulator of *Mtb*-MenD, and a possible self-regulating signal in the pathogen *Mtb*. Our initial analysis suggests that protein-level allosteric regulation of MenD by DHNA has limited direct conservation across microorganisms, which is consistent with the observation that allosteric sites are often organism-specific, being under less pressure to be stringently conserved. They may then provide species-specific sites for drug development. Our discovery of a potent inhibitor of *Mtb*-MenD, acting through a site remote from the active site may thus have important implications for future development of selective antitubercular drugs.

## Acknowledgements

This work was primarily supported by a Marsden Fund from the Royal Society Te Apārangi (M1208, to JMJ, GB and EMMB), with additional funding support from the Auckland Medical Research Foundation (to JMJ), the Canterbury Medical Research Foundation (to JMJ) and the Biomolecular Interaction Centre (to JMJ and NATH). GB is supported by a Sir Charles Hercus Fellowship through the Health Research Council of New Zealand. This research was undertaken in part using the MX2 beamline at the Australian Synchrotron, part of ANSTO, and made use of the Australian Cancer Research Foundation (ACRF) detector. Access to the Australian Synchrotron was supported by the New Zealand Synchrotron Group Ltd. We are grateful to Drs. Deborah Crittenden and Timothy Allison for helpful discussions and Dr Julia Bates for scientific editing of the manuscript.

## Author Contributions

GB initiated, supervised and designed the research, carried out the research and data analysis and contributed to manuscript construction and editing. EMMB supervised and designed the research, carried out the research and data analysis and co-wrote the manuscript. LVN and ENMJ carried out the research and contributed to data analysis. TS, NATH and SSD contributed to data analysis and contributed to manuscript construction and editing. ENB contributed to structure analysis and to manuscript structure, construction and editing. JMJ initated, supervised and designed the research, carried out the structural biology experiments, structure analysis and deposition and co-wrote the manuscript. All authors proof read the final versions of the manuscript.

The structures presented in this paper have all been deposited in the Protein Data Bank (PDB) with the following codes: 6O04, 6O0G, 6O0J, 6O0N

## Materials and Methods

### Strains and plasmids

The open reading frame encoding MenD (Rv0555) from *M. tuberculosis* H37Rv was previously cloned into the pYUB28b vector [13]. MenD mutants were generated using the pYUB28b-*menD* construct and oligonucleotide primers (**Supplementary Table 2**, Integrated DNA Technologies) with iProof^TM^ high-fidelity DNA polymerase (Bio-Rad). The PCR products were then treated with DpnI and ligated using T4 DNA ligase (Roche), before being transformed into *E. coli* TOP10 cells. The mutations were verified by DNA sequencing.

### Protein expression and purification

WT and mutant MenD constructs were expressed in *M. smegmatis* mc^2^ 4517 cells and purified using immobilized metal affinity chromatography (IMAC) and size-exclusion chromatography (SEC) as described previously [13]. In brief, cells were lysed in 20 mM HEPES pH 8.0, 150 mM NaCl, 5 mM MgCl_2_, 20 mM imidazole, 5% glycerol and 1 mM Tris(2-carboxyethyl)phosphine hydrochloride (TCEP) using a Microfluidics cell disruptor (Newton, USA). The recombinant *Mtb*-MenD protein was purified by IMAC with 5 mL HisTrap HP columns (GE Healthcare) using an imidazole gradient of 20–500 mM over 90 mL. The eluted protein solution was then concentrated and further purified by SEC (in buffer containing 20 mM HEPES pH 8.0, 150 mM NaCl, 5 mM MgCl_2_, 5% glycerol and 1 mM TCEP) using a Superdex 200 10/30 column. The protein solution was kept at −80°C for subsequent use (with added 5% glycerol).

For NMR experiments, the protein was either purified directly into 50 mM phosphate pH 7.5 with 50-100 mM NaCl or buffer-exchanged into this after purification.

### NMR spectroscopy assay

The activity of *Mtb*-MenD was monitored using a coupled reaction with *E. coli* isochorismate synthase (*Ec*-MenF), which converts chorismate to isochorismate (the substrate for *Mtb*-MenD). *Ec*-MenF was expressed and purified as previously described [13]. Initial NMR samples were prepared with 2 mM chorismate, 1 mM α-ketoglutarate, 200 µM ThDP, 25 µM *Ec*-MenF, and varying concentrations of DHNA (in 20 mM potassium phosphate pH 7.5, 50 mM NaCl, 1 mM MgCl_2_, 1 mM 2-mercaptoethanol, 10% v/v D_2_O and 0.25 mM trimethylsilyl propanoic acid). Samples were incubated at 25 ºC and 1D ^1^H NMR spectra monitored until the reaction reached equilibrium, with an estimated 47:53 ratio of isochorismate to chorismate based on peak integrals. *Mtb*-MenD (5 µM) was then added and 1D ^1^H NMR spectra were recorded at 100 s intervals for up to 90 min. Reaction rates were estimated by monitoring the decrease in peak integral for isochorismate and chorismate, and the increase in peak integrals for SEPHCHC, relative to the peak for the TSP internal standard (δ 0 ppm). Due to peak overlap in other regions, the pyruvyl methylene proton peak was monitored with a chemical shift of 5.17, 5.15 and 5.12 ppm for isochorismate, chorismate and SEPHCHC, respectively. NMR spectra were collected on an Avance AVIII-HD 500 MHz Spectrometer (Bruker) with suppression of the water signal using excitation sculpting [43]. Data were processed using the software package TopSpin 4.0.6 (Bruker).

### UV-Visible spectroscopy assay

*Mtb*-MenD activity was monitored by the decrease in isochorismate absorbance at 278 nm (ε_278_ = 8,300 M^−1^.cm^−1^; [29]) at 25 °C. To produce isochorismate, 10 µM *Ec*-MenF was incubated in a 3 mL reaction for 2 h at room temperature with 100 mM Tris HCl pH 7.5, 100 mM NaCl, 5 mM MgCl_2_ and at least 3 mg chorismic acid. *Ec*-MenF was then removed from the mixture using a vivaspin concentrator with a 10 kDa cut-off. The mixture was stored in small aliquots at −80°C prior to use.

Isochorismate was quantified prior to kinetic assays, using the following reaction: 1 µM *Mtb*-MenD was incubated with 100 µM thiamine pyrophosphate (ThDP) and 100 µM α-ketoglutarate for 30 min at 25 °C in MenD kinetic assay buffer (100 mM Tris pH 8, 100 mM NaCl and 5 mM MgCl_2_). The reaction was initiated with 30 µl of a mixture of chorismate/isochorismate. The quantity of isochorismate used was back-calculated using Beer’s law. No background catalytic rate was observed when either isochorismate or α-ketoglutarate were absent from the assay mixture. All assays were performed using a Cary 400 UV-VIS spectrophotometer and quartz cuvettes with a final reaction volume of 800 µl.

Inhibition assays for WT *Mtb-*MenD contained 0.6 µM *Mtb*-MenD, 300 µM ThDP and various concentrations of DHNA (0 to 10 µM) in a reaction buffer of 100 mM Tris pH 8, 100 mM NaCl and 5 mM MgCl_2_. After pre-incubation for 30 min at 25°C, 2 µM isochorismate was added and the reaction was initiated by the addition of 300 µM of α-ketoglutarate. Solutions of DHNA were prepared immediately prior to the inhibition assays as gradual oxidation of DHNA was observed when solutions were stored. The IC_50_ of DHNA for each mutant *Mtb*-MenD was determined using the same conditions as those for WT *Mtb-*MenD, except that the DHNA concentrations varied from 0 to 50 µM. Initial rate data were fitted to the four-parameter logistic Hill equation with GraphPad Prism.

### Crystallization, soaking and freezing

***APO_DHNA structure:*** The WT MenD apo-enzyme was concentrated to 15– 20 mg/mL in buffer A (20 mM HEPES pH 8.0, 150 mM NaCl, 5% glycerol, 1 mM TCEP, 5 mM MgCl_2_) and crystallized using 96-well sitting drop format in a variant of the MORPH crystallization screen (0.1 M MOPS/HEPES pH 7.5, 20% glycerol, 8% PEG 4K, 0.02 M CA mix; where CA mix is a mixture of sodium formate, ammonium acetate, sodium citrate tribasic dihydrate, sodium oxamate and potassium sodium tartrate tetrahydrate). Crystals grew within 1–7 days and were soaked in 10 mM DHNA solution (5% DMSO: 95% 0.1 MOPS/HEPES pH 7.5, 25% glycerol, 10% PEG 4K, 0.02 M CA mix) solution overnight before being flash-frozen in liquid nitrogen.

***ThDP, Intermediate I or II + DHNA structures:*** The WT MenD apo-enzyme was concentrated to 15–20 mg/ml in buffer A, thiamine diphosphate was added to a final concentration of 1 mM and the protein crystallized as for the apo-enzyme. Crystals grew within 1–7 days and were soaked as follows: for ThDP_DHNA, soaked in 5 mM DHNA (0.1 MOPS/HEPES pH 7.5, 25% glycerol, 14% PEG 4K, 0.01 M CA mix) for 30 min; for IntI_DHNA, soaked in 1 mM α-ketoglutarate (in 0.1 MOPS/HEPES pH 7.5, 25% glycerol, 14% PEG 4K, 0.01 M CA mix) for 1–10 min followed by addition of DHNA (final concentration 3.3 mM) to the α-ketoglutarate soak drop for a further 20 min; for Int II_DHNA, isochorismate (final concentration 100 µM) was added to an α-ketoglutarate soak drop for less than 1 min, followed by the addition of DHNA (final concentration 3.3 mM) for a further 20 min. All crystals were then flash-frozen in liquid nitrogen.

***Other soaks*** into *Mtb*-MenD crystals were undertaken using solutions of crystallization mother liquor containing varying concentrations (up to 50 mM) of either menaquinones (MK-2, MK-2 H2, MK-3 obtained from Prof. Debbie Crans [44]) or enzymatically produced (1R,6R)-2-succinyl-6-hydroxy-2,4-cyclohexadiene-1-carboxylate (SHCHC; the product of MenH) or 2-Succinylbenzoate (O-succinyl benzoic acid (OSB); the product of MenC).

### Data collection, structure determination and refinement

All diffraction data were collected using the macromolecular crystallography beamline MX2 at the Australian Synchrotron. The data were indexed and processed using XDS [45], re-indexed using POINTLESS [46] and scaled with SCALA [46] from the CCP4 program suite [47]. Analyses of merged CC_½_ correlations between intensity estimates from half data sets were used to influence high resolution cutoff for data processing [48].

The structures of the DHNA-soaked crystals were solved by molecular replacement using Phaser [49], with 5ERY [13] as a search model, and a dimer of a previous *Mtb*-MenD structure (PDB code: 5ESU [13]) was used as a search model for the ThDP/Intermediate structures. The final models of all structures were then completed with iterative rounds of manual building using COOT [50] and refinement using Refmac5 [47] and Phenix[51]. After building, additional density corresponding to DHNA, ThDP, intermediate I or II, and α-ketoglutarate, as appropriate, was modelled using available PDB dictionary restraints. Water molecules were identified by their spherical electron density and appropriate hydrogen bond geometry with the surrounding structure. Unless otherwise stated all protein structure images were generated using Pymol (The PyMOL Molecular Graphics System, Version 1.5, Schrödinger, LLC).

