## Supplementary Table 1 (ST1), Supplementary Table 2 (ST2), Supplementary Figure 1 (SF1) for "Allosteric Regulation of Vitamin K2 Biosynthesis in a Human Pathogen"

**Supplementary Table 1 (ST1).** Data collection and refinement statistics.

| | **IntII_DHNA (6O04)** | **IntI_DHNA (6O0G)** | **ThDP_DHNA (6O0J)** | **APO_DHNA (6O0N)** |
|---|---|---|---|---|
| **Data Collection** | | | | |
| X-ray source | MX2 λ = 0.95370 Å | MX2 λ = 0.95370 Å | MX2 λ = 0.95370 Å | MX2 λ = 0.95370 Å |
| Space group | P212121 | P212121 | P212121 | P212121 |
| Unit cell lengths (Å) | *a=100.80* | *a=100.67* | *a=101.55* | *a=102.18* |
| | *b=143.31* | *b=143.45* | *b=143.67* | *b=143.23* |
| | *c=173.62* | *c=172.74* | *c=176.11* | *c=184.69* |
| Resolution range (Å) [a] | 48.46–2.50 (2.54-2.50) | 48.39–2.40 (2.44-2.40) | 48.81–2.35 (2.39-2.35) | 48.12–3.03 (3.12–3.03) |
| Total reflections[a] | 908795 (46229) | 1459118 (66941) | 1598311 (79354) | 797300 (69226) |
| No. of unique reflections[a] | 87576 | 98344 | 107806 | 53451 |
| Multiplicity[a] | 10.4 (10.5) | 14.8 (14.0) | 14.8 (15.0) | 14.9 (15.2) |
| $R_{merge}$[a] | 0.361 (3.235) | 0.284 (3.150) | 0.280 (4.349) | 0.316 (4.555) |
| | (all I+ and I-) | (all I+ and I-) | (all I+ and I-) | (all I+ and I-) |
| CC ½[a] | 0.987 (0.301) | 0.996 (0.348) | 0.999 (0.356) | 0.999 (0.314) |
| $\langle I/\sigma(I)\rangle$[a] | 7.9 (0.8) | 11.6 (1.0) | 10.7 (0.7) | 10.0 (0.8) |
| Completeness[a] (%) | 100.0 (100.0) | 100.0 (99.9) | 100.0 (100.0) | 100.0 (100.0) |
| Wilson B (Å$^2$) | 42.88 | 43.44 | 46.85 | 85.65 |
| **Refinement** | | | | |
| Resolution range (Å) [a] | 48.40–2.50 (2.50–2.53) | 48.33–2.40 (2.40–2.43) | 48.79–2.35 (2.35–2.38) | 48.12–3.03 (3.03–3.09) |
| $R/R_{free}$ | 0.2045/0.2404 | 0.1925/0.2409 | 0.2055/0.2505 | 0.2170/0.2565 |
| No. of reflections (working/test) | 83151/4370 | 93373/4848 | 102250/5372 | 50712/2635 |
| No. of non-H atoms | 4329 | 16144 | 15922 | 14711 |
| No. of atoms protein/waters/ligands | 3924/382/175 | 15525/424/193 | 15304/471/147 | 14637/14/60 |
| **B factors** (Å$^2$) | | | | |
| Average all atoms | 47.8 | 52.5 | 57.3 | 78.7 |
| Protein | 48.1 | 52.7 | 57.7 | 78.7 |
| Water | 41.0 | 47.0 | 48.1 | 63.5 |
| Ligands (all/DHNA) | 48.0/37.5 | 50.5/40.4 | 46.0/41.5 | 71.8/71.8 |
| Molprobity Score | 1.42 (100th %ile) | 1.39 (100th %ile) | 1.36 (100th %ile) | 1.61 (100th %ile) |
| Est.coordinate error (Å) (max likelihood) | 0.36 | 0.34 | 0.38 | 0.47 |
| Favored/poor (%) | 88.21/0.12 | 86.96/0.19 | 88.79/0.25 | 78.48/0.00 |
| Clashscore | 5.45 (99th %ile) | 5.30 (99th %ile) | 4.71 (99th %ile) | 8.72 (97th %ile) |
| Bond lengths RMSZ | 0.003 | 0.005 | 0.005 | 0.003 |
| Bond angles RMSZ | 0.63 | 0.75 | 0.73 | 0.58 |
| Ramachandran (%) favored/allow/outliers | 97.3/2.6/0.05 | 97.6/2.4/0.05 | 97.3/2.6/0.05 | 97.2/2.8/0.05 |

[a]Values in parentheses correspond to the highest resolution shell.

**Supplementary Table 2 (ST2).** Primers used in the site-directed mutagenesis of *Mtb*-MenD constructs used in this study.

| **Construct** | | **Primer Sequences (5'-3')** |
|---|---|---|
| R97A | Forward | GCCGTGCCGCTGATCGTGCTGTCAG |
| | Reverse | AGCGTAGTTTGCCTCCACCACCG |
| R277A | Forward | GCCCCGACACTGCATCGTCCGGTATC |
| | Reverse | GCCCAGCATGATCACCTGTTGAGG |
| R303A | Forward | GCCTGGCCGGATGTCTCGGGTAACTC |
| | Reverse | TGGACCGGTTGTCAATGCGAATACCG |

### Supplementary Figure 1 (SF1).

```
                                        α1                 β1               α2                  β2          α3
Mycobacterium_tuberculosis
                                         10        20        30        40        50        60
Mycobacterium_tuberculosis              TTQARVVVDELIRGGVRDVVLCPGSRNAPLAFALQDADRSGRIRL.HVRIDERTAGYLAI
Mycobacterium_canettii                  TTQARVVVDELIRGGVRDVVLCPGSRNAPLAFALQDADRSGRIRL.HVRIDERTAGYLAI
Rhodococcus                             TAQATAVVDELVRGGVREVVLCPGSRNAPLAFALQAADLEGRLRL.HMRIDERTAGFLAL
Clostridum_paraputrificum               EMVAALIVDELIRCGTREAVVCPGSRSAPLARALLSAERAGRLRL.HVRADERSAAFLAL
Pseudomonas_aeruginosa                  PQLARAVAAELARH.LTDVVICPGSRNSPLSLELL...ARGDIRV.HTRIDERSAAFLAL
Phaeospirillum_molischianum             AAWAAALIDGLAGAGVCRAVISPGSRSTPLALACL...RHSGIVS.RVLLDERVAAFYAL
Sulfuricella_denitrificans              LRWAMALVDRLVASGVQHAVISPGSRSTPLTLACL...RHPHLRT.WIQVDERSAGFFAL
Afifella_pfennigii                      FRWAAALFAGLAAAGVRDVVVSPGSRSTPLTLAAL...RQEGLVV.RVVVDERSAGFFAL
Tropheryma_whipplei                     SGIAFCLLNLLAERGVRDIVVAPGARSQALALAAVALERAGLVRL.TVRIDERVAAFTAF
Listeria_monocytogenes                  TDYLAAFIEELVQAGVKEAIISPGSRSTPLALMMA...EHPILKI.YVDVDERSAGFFAL
Bacillus_subtilis                       THYIGSFIDEFALSGITDAVVCPGSRSTPLAILAA...AHPDISV.HVQIDERSAGFFAL
Thermosynechococcus_elongatus           VLWASVLFETLYRLGLRTVVLSPGSRSGPLAIAAA...AHPHLEA.LPILDERSAAFFAL
Clostridium_pasteurianum                TNYIAALVDELYELGVREVVISPGSRSTPLAYLFC...EH.NFNN.FVNVDERSSAFFAL
Staphylococcus_aureus                   TKQVFTFASELYAYGVREVVISPGSRSTPLALAFE...AHPNIKT.WIHPDERSAAFFAV
Halobacterium_salinarum                 TLWARAIADELAAAGVHAVCVCPGSRSTPLTVAVD...AHDDLTV.YSHLDERSAAFFAL
Cytophaga_hutchinsonii                  MQAIYDIPEILFQHGVTDVVLSPGSRCAPLSIAFS...RHKKLQI.KVVPDERSAAFIAL
Escherichia_coli                        RRWAAVILEALTRHGVRHICIAPGSRSTPLTLAAA...ENSAFIH.HTHFDERGLGHLAL
Salmonella_enterica                     RRWAAVILEALTRHGVRHVCIAPGSRSTPLTLAAA...ENPAFIH.HTHFDERGLGHLAL
Haloarcula_marismortui                  TLWAETLVGELVAGGVDAVCLSPGSRSTPLTVAFA...EHPDIEV.FSHLDERSAAFFAL
Parachlamydia_acanthamoebae             KILALSVLEEAIKKGVREFCVCAGSRNADFVKILN...QAIGVQV.YSWFEERSAGYFAL
Natronomonas_pharaonis                  TLWGQTVADELATVGIETAVLAPGSRSTPLAVAFA...QHDDIEA.VSLLDERSAAFFAL
Trichodesmium_erythraeum                TVWSSILVETLHHLGLTTAIISPGYRSTPLTFAFA...THPKIET.IPILDERSAAFFAL
Syntrophus_aciditrophicus               LLWSSLIVAELVKNGLDTFFISPGNRNAPLISALI...HEERSVK.KICVDERAAGYRAL
Desulfatibacillum_alkenivorans          VLWSSVLVEELRRNGLDAFFVSPGNRNAPLVSALA...HNDQVQK.FSCMDERGAAYRAL
Flavobacterium_psycrophilum             IPLAQSIIEICNTKGLHHIVISPGSRNAPLTIGFT...NNPVFTC.YSIADERCAAFFAL
Campylobacter_blaseri                   KKQATLLVGLLRAYDIKDVVISSGSRNGPFIHTFS...GCDFFDC.KTIVDERSAGYFAL
Candidatus_Protochlamydia_amoebophila   HELALHVIEQALQYGIKEFCVAPGARNAAFVYLLA...QIPTIKV.YYWSEERSAAFFAV
Haemophilus_influenzae                  RCWSKVILETLVRQGVSHVCIAPGSRSTPLTLEAVRLQNAGAVTC.HTHFDERGLGFFAL
Azoarcus_communis                       NKLILHLISLMKQHGILRAVISPGSRHYPLIRSME...QDPDFTL.YSVVDERSAAFFAL
Prochlorococcus_marinus                 FFISLELLKTLVAKGVKYFVLCPGSRSGPLALAAASLSKRKELTL.ITSIDERSAAFLAL
Neochlamydia_sp.                        EKLVIQVLQELIYRGVKEICICPGGRNVPFITALE...KAEQFKL.YYWPEERSAAFFAL
Arabidopsis_thaliana                    AVWASAIIEECTRLGLTYFCVAPGSRSSHLAIAAA...NHPLTTC.LACFDERSLAFHAI
Campylobacter_insulaenigrae             KKSIQLLAHHLDALGVHHFVISPGSRNAPIAIHFT...QIENFHC.YSIVDERSAAFVAM
Gramella_forsetii                       IPVARSVVALCVAKDIKHVVISPGSRNAPLTIGFT...HHDEITP.YSIVDERCAAFFAL
Burkholderia_sp.                        AVWVECIVEMLSAAGVRHAVLCPGGRASAMCLAVD...AHPRIAVEMVCTDERSGAFVAL
```

```
                                                  β3     α4      α5            β4        η1
Mycobacterium_tuberculosis
                                              70   ★    80        90       100       110       120
Mycobacterium_tuberculosis              GLAIGAGAPVCVAMTSGTAVANLGPAVVEANYARVPLIVLSANRPYELLGTGANQTMEQL
Mycobacterium_canettii                  GLAIGAGAPVCVAMTSGTAVANLGPAVVEANYARVPLIVLSANRPYELLGTGANQTMEQL
Rhodococcus                             GLAVAGKRPVPIVMTSGTAVANLGPAVLEANYARVPLVVLSANRPYEMLGTGANQTIEQL
Clostridum_paraputrificum               GLAAATGTVTPVAVTSGTAVANLLPAMVEATYSGVPLMALTADRPASYRGTGANQTIVQD
Pseudomonas_aeruginosa                  GMGRASGRHVGIVTTSGTAVANCLPAIVEAYHSHTPIAMISADRPARLQGTGANQTINQR
Phaeospirillum_molischianum             GLARAEEKPVVLVCTSGSAAANWYPAVAEADAARVPLILLTADRPPELQDCGANQTFDQR
Sulfuricella_denitrificans              GLSKADRRPVVLICTSGSAPANWFPAVIEANHGLTPLILLTADRPPELRDCGANQTIDQV
Afifella_pfennigii                      GIAKSTGHPVALVATSGSAVANWMPAVVEANMGEMPLILLSADRPPELQDCGANQTMNQI
Tropheryma_whipplei                     GMAKALDSPAVVITTSGSAVANLYPAVVEASHSFVPLIVITADRPRELHGKRSNQTMNQE
Listeria_monocytogenes                  GLAKASKRPVVLLCTSGTAAANYFPAVAEANLSQIPLIVLTADRPHELRNVGAPQAMDQL
Bacillus_subtilis                       GLAKAKQRPVLLICTSGTAAANFYPAVIEAHYSRVPIIVLTADRPHELREVGAPQAINQH
Thermosynechococcus_elongatus           GLVQQQGRPVALLCTSGTAAANFYPAIIEASLSHLPLIVLSADRPPELRFCQAGQAIDQV
Clostridium_pasteurianum                GIAKEHERPVVLVCTSGSAAANYLPAIVEAKYSGVPLIILTADRPPELRNVGAPQTIDQN
Staphylococcus_aureus                   GLIKGSERPVAILCTSGTAAANYTPAIAESQISRIPLIVLTSDRPHELRSVGAPQAINQV
Halobacterium_salinarum                 GRGKRTGAPTAVLTTSGTATANLHPAVMEAAQARIPLVVLTADRPPELRGSGANQTVDQE
Cytophaga_hutchinsonii                  GMSLQTQKPTVLICTSGSALYNYAPAIVEAYYQRVPLIVLSADRPPEWIDQNDGQTIRQQ
Escherichia_coli                        GLAKVSKQPVAVIVTSGTAVANLYPALIEAGLTGEKLILLTADRPPELIDCGANQAIRQP
Salmonella_enterica                     GLAKVSQQPVAVIVTSGTAVANLYPALIEAGLTGEKLILLTADRPPELIDCGANQAIRQA
Haloarcula_marismortui                  GRARRTGEPTPLVCTSGTAAANYHPAVIEANQSGVPLLLLTADRPPELIDSGANQTVDQE
Parachlamydia_acanthamoebae             GRSKQTGRPVGVITTSGTAAGELLPAMMEGYYSGLPLLAITADRPRRLRGTGAPQTAEQV
Natronomonas_pharaonis                  GYAKRTGQPAPLVCTSGTALANFHPAVIEADTARVPMVLLTADRPPELADSGANQTIDQA
Trichodesmium_erythraeum                GIAKKNYQPIVIVCTSGTAAANFYPAIIEAKESRIPLLVLTADRPPELRNCHAGQTIDQV
Syntrophus_aciditrophicus               GHAKAAGRPGVLVCTSGTAPANYYPAVIEAFRDEIPLVILSADRPPELIGSDANQTIVQP
Desulfatibacillum_alkenivorans          GYAKATGKPGVLVCTSGTALANYHPAVIEAWRDELPLMVVSADRPSELTGSDANQTISQE
Flavobacterium_psycrophilum             GIAQQLKKPVALVCTSGSALLNYYPAFAEAFYSQIPLVVISADRPQSKIDIGDGQTIRQE
Campylobacter_blaseri                   GLARASKKPVVLVCSSGTATLNYAPSIAEAYYQKVPLIVITADRPNYYISQMENQCLNQD
Candidatus_Protochlamydia_amoebophila   GRIKATHQPIAVVTTSGTAVAELLPATIEAYYTSLPLLLITADRPRRFRLSGAPQSIEQV
Haemophilus_influenzae                  GIAKATQSPVAIIVTSGTATANLYPAIIEARQTGVNLFVLTADRPPELWECGANQAILQQ
Azoarcus_communis                       GLIQQTGAPVAVCCTSGTSAINYGSAVVEAFYQGLPLVVLTADRLPELLNQKEEQMCLQK
Prochlorococcus_marinus                 GISAASGQVSCVITTSGSAVANLLPAAVEADRSCHPLLFLTADRPLRLKECGANQAVNQQ
Neochlamydia_sp.                        GRSKLMHRPTAVLTTSGTAVGELLPAAMEAYYTGIPLILLTADRPRRFRRTGSPQTAEQV
Arabidopsis_thaliana                    GYAKGSLKPAVIITSSGTAVSNLLPAVVEASEDFLPLLLLTADRPPELQGVGANQAINQI
Campylobacter_insulaenigrae             GMAKRLGKPVGVICTSGSAAVNYYPAVTEAFYQNIPLLLLTADRPTDFVDLFDGQTIRQK
Gramella_forsetii                       GLAQELKKPVALVCTSGSALLNYYPAIAEAYYSDIPLVIISADRPIERIDIGDGQTIRQK
Burkholderia_sp.                        GIAKASGSPVAIVTTSGSAVANVLPALTEADAADVPLVVLTCDRPRALRGTGFGQMADHL
```

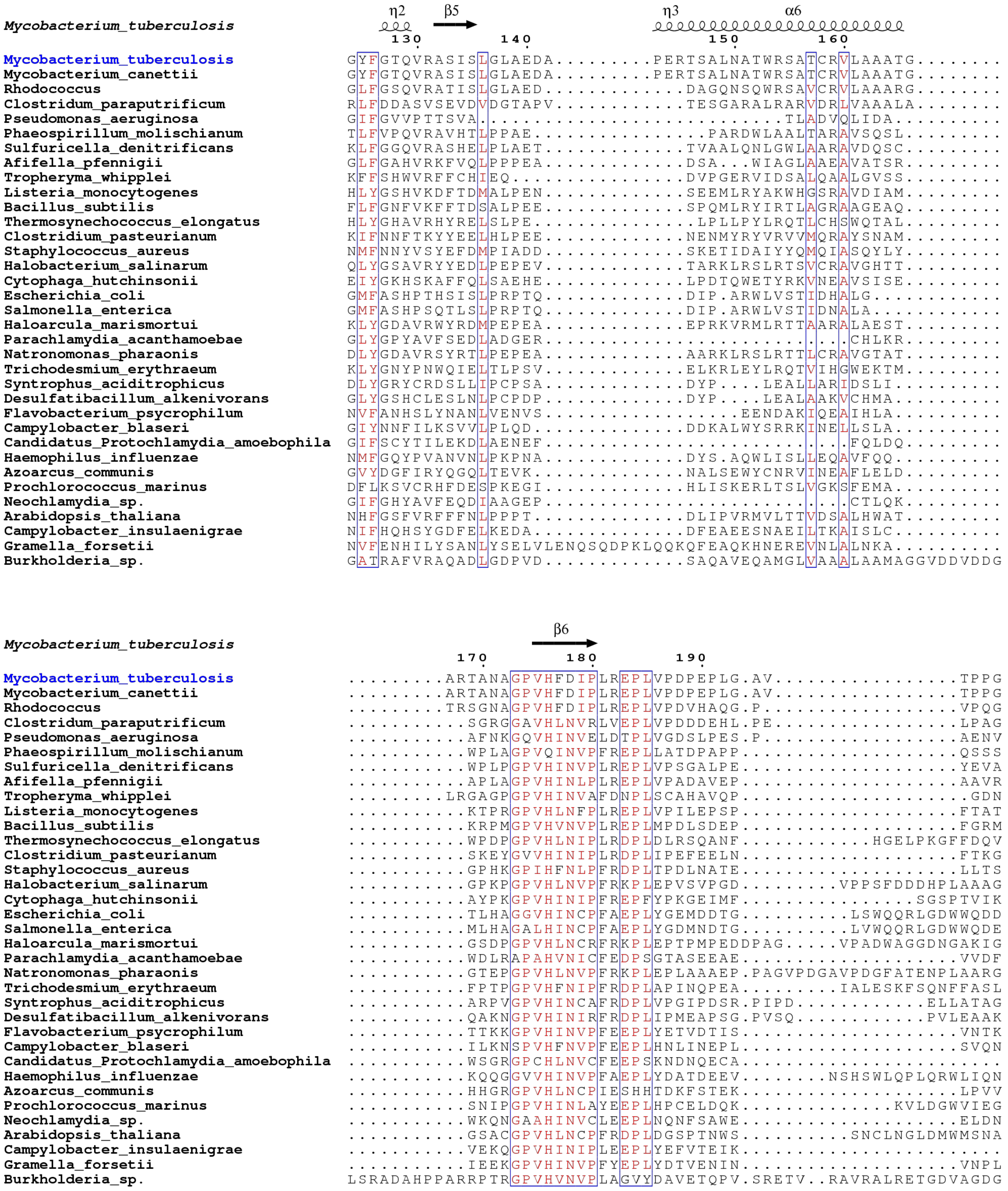
Mycobacterium_tuberculosis
η2 β5 η3 α6
130 140 150 160
Mycobacterium_tuberculosis GYFGTQVRASISLGLAEDA.........PERTSALNATWRSATCRVLAAATG........
Mycobacterium_canettii GYFGTQVRASISLGLAEDA.........PERTSALNATWRSATCRVLAAATG........
Rhodococcus GLFGSQVRATISLGLAED............DAGQNSQWRSAVCRVLAAARG........
Clostridum_paraputrificum RLFDDASVSEVDVDGTAPV...........TESGARALRARVDRLVAAALA.......
Pseudomonas_aeruginosa GIFGVVPTTSVA.......................TLADVQLIDA.........
Phaeospirillum_molischianum TLFVPQVRAVHTLPPAE..............PARDWLAALTARAVSQSL........
Sulfuricella_denitrificans KLFGGQVRASHELPLAET............TVAALQNLGWLAARAVDQSC........
Afifella_pfennigii GLFGAHVRKFVQLPPPEA.............DSA...WIAGLAAEAVATSR........
Tropheryma_whipplei KFFSHWVRFFCHIEQ..............DVPGERVIDSALQAALGVSS........
Listeria_monocytogenes HLYGSHVKDFTDMALPEN.............SEEMLRYAKWHGSRAVDIAM.......
Bacillus_subtilis FLFGNFVKKFFTDSALPEE............SPQMLRYIRTLAGRAAGEAQ........
Thermosynechococcus_elongatus HLYGHAVRHYRELSLPE..............LPLLPYLRQTLCHSWQTAL........
Clostridium_pasteurianum KIFNNFTKYYEELHLPEE.............NENMYRYVRVVMQRAYSNAM.......
Staphylococcus_aureus NMFNNYVSYEFDMPIADD............SKETIDAIYYQMQIASQYLY........
Halobacterium_salinarum QLYGSAVRYYEDLPEPEV............TARKLRSLRTSVCRAVGHTT........
Cytophaga_hutchinsonii EIYGKHSKAFFQLSAEHE............LPDTQWETYRKVNEAVSISE........
Escherichia_coli GMFASHPTHSISLPRPTQ............DIP.ARWLVSTIDHALG..........
Salmonella_enterica GMFASHPSQTLSLPRPTQ............DIP.ARWLVSTIDNALA..........
Haloarcula_marismortui KLYGDAVRWYRDMPEPEA............EPRKVRMLRTTAARALAEST........
Parachlamydia_acanthamoebae GLYGPYAVFSEDLADGER......................CHLKR........
Natronomonas_pharaonis DLYGDAVRSYRTLPEPEA............AARKLRSLRTTLCRAVGTAT........
Trichodesmium_erythraeum KLYGNYPNWQIELTLPSV............ELKRLEYLRQTVIHGWEKTM........
Syntrophus_aciditrophicus DLYGRYCRDSLLIPCPSA............DYP....LEALLARIDSLI.........
Desulfatibacillum_alkenivorans GLYGSHCLESLNLPCPDP............DYP....LEALAAKVCHMA.........
Flavobacterium_psycrophilum NVFANHSLYNANLVENVS................EENDAKIQEAIHLA.........
Campylobacter_blaseri GIYNNFILKSVVLPLQD.............DDKALWYSRRKINELLSLA.........
Candidatus_Protochlamydia_amoebophila GIFSCYTILEKDLAENEF.......................FQLDQ........
Haemophilus_influenzae NMFGQYPVANVNLPKPNA............DYS.AQWLISLLEQAVFQQ.........
Azoarcus_communis GVYDGFIRYQGQLTEVK.............NALSEWYCNRVINEAFLELD........
Prochlorococcus_marinus DFLKSVCRHFDESPKEGI............HLISKERLTSLVGKSFEMA.........
Neochlamydia_sp. GIFGHYAVFEQDIAAGEP......................CTLQK........
Arabidopsis_thaliana NHFGSFVRFFFNLPPPT.............DLIPVRMVLTTVDSALHWAT........
Campylobacter_insulaenigrae NIFHQHSYGDFELKEDA.............DFEAEESNAEILTKAISLC........
Gramella_forsetii NVFENHILYSANLYSELVLENQSQDPKLQQKQFEAQKHNEREVNLALNKA..........
Burkholderia_sp. GATRAFVRAQADLGDPVD............SAQAVEQAMGLVAAALAAMAGGVDDVDDG
Mycobacterium_tuberculosis
β6
170 180 190
Mycobacterium_tuberculosis .........ARTANAGPVHFDIPLREPLVPDPEPLG.AV................TPPG
Mycobacterium_canettii .........ARTANAGPVHFDIPLREPLVPDPEPLG.AV................TPPG
Rhodococcus .........TRSGNAGPVHFDIPLREPLVPDVHAQG.P.................VPQG
Clostridum_paraputrificum ..........SGRGGAVHLNVRLVEPLVPDDDEHL.PE................LPAG
Pseudomonas_aeruginosa ...........AFNKGQVHINVELDTPLVGDSLPES.P.................AENV
Phaeospirillum_molischianum ..........WPLAGPVQINVPFREPLLATDPAPP...................QSSS
Sulfuricella_denitrificans ..........WPLPGPVHINVPLREPLVPSGALPE...................YEVA
Afifella_pfennigii ..........APLAGPVHINLPLRDPLVPADAVEP...................AAVR
Tropheryma_whipplei ........LRGAGPGPVHINVAFDNPLSCAHAVQP....................GDN
Listeria_monocytogenes ..........KTPRGPVHLNFPLREPLVPILEPSP...................FTAT
Bacillus_subtilis ..........KRPMGPVHVNVPLREPLMPDLSDEP...................FGRM
Thermosynechococcus_elongatus ..........WPDPGPVHLNIPLRDPLDLRSQANF...........HGELPKGFFDQV
Clostridium_pasteurianum ..........SKEYGVVHINIPLRDPLIPEFEELN...................FTKG
Staphylococcus_aureus ..........GPHKGPIHFNLPFRDPLTPDLNATE...................LLTS
Halobacterium_salinarum ..........GPKPGPVHLNVPFRKPLEPVSVPGD........VPPSFDDDHPLAAAG
Cytophaga_hutchinsonii ..........AYPKGPVHINIPFREPFYPKGEIMF...............SGSPTVIK
Escherichia_coli ..........TLHAGGVHINCPFAEPLYGEMDDTG.........LSWQQRLGDWWQDD
Salmonella_enterica ..........MLHAGALHINCPFAEPLYGDMNDTG.........LVWQQRLGDWWQDE
Haloarcula_marismortui ..........GSDPGPVHLNCRFRKPLEPTPMPEDDPAG.....VPADWAGGDNGAKIG
Parachlamydia_acanthamoebae ..........WDLRAPAHVNICFEDPSGTASEEAE...................VVDF
Natronomonas_pharaonis ..........GTEPGPVHLNVPFRKPLEPLAAAEP.PAGVPDGAVPDGFATENPLAARG
Trichodesmium_erythraeum ..........FPTPGPVHFNIPFRDPLAPINQPEA........IALESKFSQNFFASL
Syntrophus_aciditrophicus ..........ARPVGPVHINCAFRDPLVPGIPDSR.PIPD...........ELLATAG
Desulfatibacillum_alkenivorans ..........QAKNGPVHINIRFRDPLIPMEAPSG.PVSQ.............PVLEAAK
Flavobacterium_psycrophilum ..........TTKKGPVHINVPFEEPLYETVDTIS...................VNTK
Campylobacter_blaseri ..........ILKNSPVHFNVPFEEPLHNLINEPL...................SVQN
Candidatus_Protochlamydia_amoebophila ..........WSGRGPCHLNVCFEEPSKNDNQECA....................
Haemophilus_influenzae ..........KQQGGVVHINVPFAEPLYDATDEEV.......NSHSWLQPLQRWLIQN
Azoarcus_communis ..........HHGRGPVHLNCPIESHHTDKFSTEK...................LPVV
Prochlorococcus_marinus ..........SNIPGPVHINLAYEEPLHPCELDQK.............KVLDGWVIEG
Neochlamydia_sp. ..........WKQNGAAHINVCLEEPLNQNFSAWE...................ELDN
Arabidopsis_thaliana ..........GSACGPVHLNCPFRDPLDGSPTNWS..........SNCLNGLDMWMSNA
Campylobacter_insulaenigrae ..........VEKQGPVHINIPLEEPLYEFVTEIK....................
Gramella_forsetii ..........IEEKGPVHINVPFYEPLYDTVENIN...................VNPL
Burkholderia_sp. LSRADAHPPARRPTRGPVHVNVPLAGVYDAVETQPV.SRETV..RAVRALRETGDVAGDG

```
                                                                                     η5
Mycobacterium_tuberculosis                 ϱ...........................................................
                                         200

Mycobacterium_tuberculosis                 RP..........................................................
Mycobacterium_canettii                     RP..........................................................
Rhodococcus                                RP..........................................................
Clostridum_paraputrificum                  R...........................................................
Pseudomonas_aeruginosa                     K...........................................................
Phaeospirillum_molischianum                L...........................................................
Sulfuricella_denitrificans                 A...........................................................
Afifella_pfennigii                         A...........................................................
Tropheryma_whipplei                        K...........................................................
Listeria_monocytogenes                     G.................KKHH......................................
Bacillus_subtilis                          R.................TGRH......................................
Thermosynechococcus_elongatus              Q...........................................................
Clostridium_pasteurianum                   R...........................................................
Staphylococcus_aureus                      E...........................................................
Halobacterium_salinarum                    RG...............GDTPF......................................
Cytophaga_hutchinsonii                     R...........................................................
Escherichia_coli                           K...........................................................
Salmonella_enterica                        K...........................................................
Haloarcula_marismortui                     R.................DGPF......................................
Parachlamydia_acanthamoebae                RH...............YEE........................................
Natronomonas_pharaonis                     R.................DGPF......................................
Trichodesmium_erythraeum                   R...........................................................
Syntrophus_aciditrophicus                  RL...............YAREG......................................
Desulfatibacillum_alkenivorans             RY...............FKNTA......................................
Flavobacterium_psycrophilum                I...........................................................
Campylobacter_blaseri                      R...........................................................
Candidatus_Protochlamydia_amoebophila      R...........................................................
Haemophilus_influenzae                     K...........................................................
Azoarcus_communis                          R...........................................................
Prochlorococcus_marinus                    F...........................................................
Neochlamydia_sp.                           R...........................................................
Arabidopsis_thaliana                       EP...............FTKYF......................................
Campylobacter_insulaenigrae                ............................................................
Gramella_forsetii                          Q...........................................................
Burkholderia_sp.                           RGMAMRRDVGGGAETLAFEAKAARAVERLDGHANEHVNQHANEHANEHTSEHTSEHTGER

                                                                              β7         β8
Mycobacterium_tuberculosis                 .........................ϱϱ        ━━→ ━━━..........→    ━
                                                                    210                 220

Mycobacterium_tuberculosis                 .........................AGKPWTYTPPVTFDQPLD..........IDLSVDT
Mycobacterium_canettii                     .........................AGKPWTYTPPVTFDQPLD..........IDLSVDT
Rhodococcus                                .........................GGAAWTTTQHATLDVPMD..........LDLTPDT
Clostridum_paraputrificum                  .........................GDGPWTELRHLAAGKAGNVGVAK.....LDLGKKT
Pseudomonas_aeruginosa                     .................................KAPIKSEVVDHGEVE.....IDLSRNT
Phaeospirillum_molischianum                ...........................PPLVLQPRLMLPQDSVEILAR.....RVEGRRG
Sulfuricella_denitrificans                 ..........................KPKTVSYPAMLPPADEISRWAT.....ELSGRPG
Afifella_pfennigii                         ...........................PVRLVGSCAPNEETLASLAGI.....LAAGSRG
Tropheryma_whipplei                        .............................GSALKHKKVLRSKRETLVL.....RKDDPPT
Listeria_monocytogenes                     ...........................HVHIYYTHEVLDDSSIQKMVT.....ECTGKKG
Bacillus_subtilis                          ...........................VSVKTGTQSVDRDSLSDVAAM.....LAEAEKG
Thermosynechococcus_elongatus              .............................PFVPPRVVTSLPWQT.........WQQMQRG
Clostridium_pasteurianum                   .............................FENKFEYIKGENQVIFESS.....ILKNKNG
Staphylococcus_aureus                      ..........................MKILPHYQKSIDALAIRH.........ILNKKKG
Halobacterium_salinarum                    ...........................VSVHDGTTEPAGETAAALAAA.....ATTAARP
Cytophaga_hutchinsonii                     ..............................EQPLHTLSDEQWKSIQHK.....LDSYKKI
Escherichia_coli                           ..............................PWLREAPRLESEKQRDWF.....FWRQKRG
Salmonella_enterica                        ..............................PWLREARRLESDKQRDWF.....FWRQKRG
Haloarcula_marismortui                     ...........................VTTSEGVETPDEQTVRRVQDA.....LEAAERG
Parachlamydia_acanthamoebae                ..........................ELLQASYLPENPL.NRALLDQF.....FMVSKKP
Natronomonas_pharaonis                     ...........................VEVHSGCTDPSATTVDELATA.....VEAAASG
Trichodesmium_erythraeum                   .............................PIIRTELIPNSDLIELLKN.....QFKSHSG
Syntrophus_aciditrophicus                  ...........................AYTTYPSPGTLHTGLEDVEAI.....LNRTARG
Desulfatibacillum_alkenivorans             ...........................PATVYPLVRTSCPDLSAVESI.....LKKAQRG
Flavobacterium_psycrophilum                .............................IDFDTENKPLENLDVFVNR.....WNDAKKK
Campylobacter_blaseri                      ...........................VNLARIDAKLDENELAKFRDN.....LNKSSKI
Candidatus_Protochlamydia_amoebophila      ..........................SFSMGIFTPKPILCDSHLLDEF.....IQKCQFP
Haemophilus_influenzae                     .............................SWINVEAQQNEVLMHENWD.....HWRTKRG
Azoarcus_communis                          .............................RITRVDADADDYVWQRYAS.....QLSGRKV
Prochlorococcus_marinus                    .............................LKGKITPTKDEVVKSFQSLK.LLKLDPFSLG
Neochlamydia_sp.                           ...........................CTKVAHQKGDIEAAKAALDQF.....VQRNKYP
Arabidopsis_thaliana                       ...........................QVQSHKSDGVTTGQITEILQV.....IKEAKKG
Campylobacter_insulaenigrae                .........................KFRTPEAMIAQSQYALPSKLATE.....WNAAKKI
Gramella_forsetii                          ...........................IFPEIKERHYSEKQLQNYANE.....WNRAERK
Burkholderia_sp.                           LNEHTNNRLSEHTNERLNEHTDERLNERVVEHLPEQIVARVMTRVLAKRGDRPLSEGLDG
```

```
                                           β9                        η6     β10
Mycobacterium_tuberculosis                                                     TT
                                            230       240       250       260

Mycobacterium_tuberculosis                 VVISGHGAG..............VHPNLAALPTVAEPTAPRSGDNPL............H
Mycobacterium_canettii                     VVISGHGAG..............VHPNLAALPTVAEPTAPRSGDNPL............H
Rhodococcus                                IVISGHGSA..............LRPELAGLPTVAEPTAPLHG.IPV............H
Clostridum_paraputrificum                  LVIAGSGAP..............DLPALADVPTIAEPNAPAPE.TPV............H
Pseudomonas_aeruginosa                     LVIAGDEAW..............DVPGLEDVPTIAEPSAPAPY.HPV............H
Phaeospirillum_molischianum                VILAGAEHV........PAEPLIRLANALNWPIIADPLSGLRF.GAH.D..........R
Sulfuricella_denitrificans                 LIVCGESEY.S....DGFPAALAQLALQLACPVLADPLSNLRF.GVH.D..........R
Afifella_pfennigii                         AIVCGSRPL.S....PPASQAVLELARRLNVPVFADILSGLRF.GSD............E
Tropheryma_whipplei                        VIIAGTGSG..........RRPADIACATGWPLVAEISSGSRF.GPF.............
Listeria_monocytogenes                     VFVVGPIDK.K.....ELEQPMVDLAKKLGWPILADPLSGLRS.YGA...........LD
Bacillus_subtilis                          MIVCGELHS.D.....TEKEHIIALSKALQYPILADPLSNLRN.GAH.D..........K
Thermosynechococcus_elongatus              LIIAGPSHGVD...PLAEAAAIDRLSRFLQWPVLADALSSARG.LPH.............
Clostridium_pasteurianum                   IIICGGDAY.S.....NYHKEVIKLGERLKVPVLADPISNFRN.YS.............K
Staphylococcus_aureus                      LIIVGDMQH.......QEVEQILTYSTIYDLPILADPLSHLRK...FD...........H
Halobacterium_salinarum                    LVVAGPADG.G...AGITPDAAAALADATGAPIFADPLSGLRF.GPHVG..........D
Cytophaga_hutchinsonii                     LFVGGQHLY.D........ESLRLKIGNIKAPFIGEVVSNLHG................V
Escherichia_coli                           VVVAGRMSA.......EEGKKVALWAQTLGWPLIGDVLSQTGQ.................
Salmonella_enterica                        VVVAGRMSA.......EEGKKVAQWAQTLGWPLIGDVLSQTGQ.................
Haloarcula_marismortui                     LIVAGPADQ......GLSADSLERLAAATGFPVLADPLSDLRF.GPHVDR........LD
Parachlamydia_acanthamoebae                LVVVSALKT.......VDQDVVVDFLVQLGAPCYLEGISGLRE.NPK...........LQ
Natronomonas_pharaonis                     LIVCGPTDR.P....APDAESLVALADATGFSVFADPLSGLRF.GPHVD..........D
Trichodesmium_erythraeum                   IIIAGLAQP.E..KPEVYCQAIAKISQTLNFPVLAEGLSPLRN.YSQ...........LN
Syntrophus_aciditrophicus                  LIVIGRLDG.P.....RDAPALEELAKKLGWPVFCDIASSMKG.RIP............S
Desulfatibacillum_alkenivorans             LLVIGRLEN.N.....DDRQGAAALASMLGWPVFCDIASSLKG..............AVG
Flavobacterium_psycrophilum                LILIGGCDP.N....VIQQFIIDFLANDTSVVVMTEVTSNVHH.................
Campylobacter_blaseri                      LIIAGQNLQ.E.....KSFESLKEFAKNTNAIILGEHLANLEK................N
Candidatus_Protochlamydia_amoebophila      LVIVGALNQ.......NETTPILSFLRHLKAPVYLEAQSGLRE.NSL...........LE
Haemophilus_influenzae                     VVVVGQLPA.......EQAMGINSWASAMGWVLLTDIQSGVVP.................
Azoarcus_communis                          LIVWGQAAPVD....ERLGQAVERFVEMFDCAILVDRLSNCRS.................
Prochlorococcus_marinus                    IIIVGPWRG.KVKQLNSFRGALKKWQKLTGWPILADPLSGVEN....D...........Q
Neochlamydia_sp.                           FVVVGGLSC.......QARQATVKFLADYQAPVYLEAASGLRE.NPY...........LR
Arabidopsis_thaliana                       LLLIGAIHT.E.....DEIWASLLLAKELMWPVVADVLSGVRL.RKLFKPFV....EKLT
Campylobacter_insulaenigrae                LILTGTLKD.N....AELQMLLAQIVKNHSAVVLTEVNSNLHD.................
Gramella_forsetii                          MVIVGVAQP.N....AVEQKFLEGLATDPSVIVLTETTSNLHQ.................
Burkholderia_sp.                           LIVVGPEPG.......VPLEAIFALAASSGFPVLADAGSGLRS.GPSALAPQTAARGAAG
```

```
                                                                   η7     β11          α7          β12
Mycobacterium_tuberculosis
                                                                    270       280       290

Mycobacterium_tuberculosis                 PLA.....................LPLLRPQQVIM.LGRPTLHRPVSVLLADAE.VPVFA
Mycobacterium_canettii                     PLA.....................LPLLRPQQVIM.LGRPTLHRPVSVLLADAE.VPVFA
Rhodococcus                                PMA.....................LPQLKPRQAVI.TGRPTLHRSVSKVLADPS.VAVYA
Clostridum_paraputrificum                  PLAAATFA................RDDCRPEQIVV.LGRPTLHRGVARLLADPR.IDTVV
Pseudomonas_aeruginosa                     PLAAGIFAHEQVSAEGY.......VVNTKPEQIVV.VGHPTLHRSVFDLIADPA.IEVTV
Phaeospirillum_molischianum                KRVMTRGDLFLR............GDFPPAEIVLR.FGAFPVSKAIGQWQSRA..IERIV
Sulfuricella_denitrificans                 SHIFSRYDAFLR.CTGF.......AGTHSPEWVLR.FGTMPVSRQLQNTLAACTGATHFL
Afifella_pfennigii                         ARLLRHPDQVAR.............AAPVPDWILR.FGGAPVSKAVQTWLENARGCTQIV
Tropheryma_whipplei                        ..LIKNYRNWLT............ENSLDIRRAIV.FGHPTLSSEIPTFLTKRN.IDTIV
Listeria_monocytogenes                     EVVIDQYDAFLK.EAEI.......MDKLTPEVVIR.FGSMPVSKPLKNWLEQLSDIRFYV
Bacillus_subtilis                          STVIDAYDSFLK.DDEL.......KAKLRPDVVIR.FGPMPVSKPVFLWLKDDPAIQQIV
Thermosynechococcus_elongatus              ..GISHYDLLLR.DAHL.......REYLRPEAVIQ.LGPLPTSKALREWLSACD.PLIWC
Clostridium_pasteurianum                   DIIVDSYDAFLK.SDDI.......KRELKPEFIIH.FGQVAVSKRLQQFLSMHKDALYLQ
Staphylococcus_aureus                      PNVICTYDLLFR.S..........GLDLKVDFVIR.VGKPVISKKLNQWLKKTD.AFQIL
Halobacterium_salinarum                    APVVGGYDGFLA............ADVPHPEFVLR.FGASPTSKPLRKWLAASD.ARQVV
Cytophaga_hutchinsonii                     RNVIHTHDTLLT.SIPLTD.....LEELKPDLVIS.FGGALLSKWLKQFIRSNESIDHWY
Escherichia_coli                           ..PLPCADLWLG.NAKAT......SELQQAQIVVQ.LGSSLTGKRLLQWQASCEPEEYWI
Salmonella_enterica                        ..PLPCADLWLG.NAKAV......TELQQAQIVVQ.LGSSLTGKRLLQWQATCEPEEYWV
Haloarcula_marismortui                     VPVCGGYDGYLG.SDSV.......DQWDDPDVVVR.FGASPTSKPLRHYLRDAD.CQQFL
Parachlamydia_acanthamoebae                HLRITRTERLWK.YAE........KADYEIDGILR.IGGIPTIRLWRDLEDQKQHVHVCS
Natronomonas_pharaonis                     APVCGGYDAYLP............ALEQTPEVVIR.FGASPTSKPLRQYLRDAD.ARQFI
Trichodesmium_erythraeum                   PYLISTYDLILR.NQKL.......ANKLIPKIVLQ.IGELPTSKQLRTWLEAAN.SHRLI
Syntrophus_aciditrophicus                  DRQIFSLDHPEA..LRL.......VSAYAPETILQ.FGSGLVSKHYFASLLPHSEATVIQ
Desulfatibacillum_alkenivorans             DREIFSLDHPKA..LAL.......VNGYAPEVILQ.LGTGLVSKHYYAAILKGGAKYIIQ
Flavobacterium_psycrophilum                ANFITNIDAIIT.PFTDDD.....FTNFQPEILIT.MGGMIVSKRIKAFLRKYKPQQHWH
Campylobacter_blaseri                      DNTIWQVDLVVA.KILNTT.....PNEFKPDLLIT.FEGQVISKNIKKYLKTFKPKFHYH
Candidatus_Protochlamydia_amoebophila      NLRIESVEDVWN.CSS........QHGYPIDGILR.IGGIPTARLWRDLEETNN.VKVCS
Haemophilus_influenzae                     ..TTPYEDIWLA.NQTVR......EKLLQADIVIQ.FGARFISKRINQFLQAFK.GEFWL
Azoarcus_communis                          SRAIKSAYLALR.AMTIDQ.....ASELAPDIVITLFGNYTFNSGLKGYLSTLD.NPFEV
Prochlorococcus_marinus                    EGLINHWDLFFS..IGL.......FEKIKEIQVLR.LGPIPPSRELQTWLKKPGKFQLLI
Neochlamydia_sp.                           TLSITRGDNIWK.AAE........KAGYPIDGILR.IGSIPTLRLWRDLEELKGRINVCS
Arabidopsis_thaliana                       HVFVDHLDHALF.SDSV.......RNLIEFDVVIQ.VGSRITSKRVSQMLEKCFPFAYIL
Campylobacter_insulaenigrae                DKFFNHIDRYIF.NFSEED.....YRAYAPDLLIT.VGQNVVSKKVKQFLRKAQPTQHWH
Gramella_forsetii                          EQFFTRIDTLIG.PIEKDENREELFNRLQPDILLT.FGGMIVSKKIKSFLRNYSPQHHWH
Burkholderia_sp.                           ALIVNAFDVFGG.AAR........LAATRAELIVR.FGLAPVMPVLHAYLEAQADVPTIK
```

```
                                                  β13
Mycobacterium_tuberculosis
                                300       310

Mycobacterium_tuberculosis            LT.TGPRWPDVSGNSQAT......................................GTRA
Mycobacterium_canettii                LT.TGPRWPDVSGNSQAT......................................GTRA
Rhodococcus                           LT.TGPRWPDVSGNVLAT......................................GTRA
Clostridum_paraputrificum             VA.AGDDYPDVSANARRV......................................VAAV
Pseudomonas_aeruginosa                LT.RTDTVTDPTRSAKKV......................................GTRV
Phaeospirillum_molischianum           VS.EDSRWPDPGRNAAMM......................................IHAD
Sulfuricella_denitrificans            VE.AYGRWPDPLHLTTRL......................................LRAD
Afifella_pfennigii                    VS.ASPRLADPGRNASHL......................................VTAD
Tropheryma_whipplei                   VAPFGREFYNPSGTARPVAN....................................VLVD
Listeria_monocytogenes                VD.PGAAWKDPIKAVTDM......................................IHCD
Bacillus_subtilis                     ID.EDGGWRDPTQASAHM......................................IHCN
Thermosynechococcus_elongatus         LD.PTGDNNNPLHGRCQM......................................LAIA
Clostridium_pasteurianum              VS.ETFYYHNPALSTKKY......................................IVSS
Staphylococcus_aureus                 VQNNDKIDVFPIAPDISY.......................................EIS
Halobacterium_salinarum               VD.PAGGWREAEFTATDV......................................VAAD
Cytophaga_hutchinsonii                VG.PDVTTPDVFQHLCQI......................................IPVQ
Escherichia_coli                      VD.DIEGRLDPAHHRGRR......................................LIAN
Salmonella_enterica                   ID.NIEGRLDPAHHRGRR......................................LVAK
Haloarcula_marismortui                VD.PAGGWSEAEFTATNL......................................LVAN
Parachlamydia_acanthamoebae           IS......ELPFSGLNRGVH....................................IQTS
Natronomonas_pharaonis                VD.PAGGWREATFTATDL......................................VVAD
Trichodesmium_erythraeum              ID.QSDHNFDPLHGKTTH......................................LRIS
Syntrophus_aciditrophicus             IS.PRAGLRDPAHRVNVR......................................LSMP
Desulfatibacillum_alkenivorans        VS.PRREHRDPAFKVNIR......................................IKAG
Flavobacterium_psycrophilum           ID..ELRAYDTFGSLTKH......................................FEVL
Campylobacter_blaseri                 ISNDDNNFIDTYEALSDV......................................LPLS
Candidatus_Protochlamydia_amoebophila IS......ENPFAGLTNGNI....................................IHTS
Haemophilus_influenzae                VE.QSGKALDPYHHSLTR......................................FNAK
Azoarcus_communis                     WDLGTSRVADPFRRLTTM......................................FAAR
Prochlorococcus_marinus               TE.GDSRNLDPIGGSTQFSE...................................GFSCW
Neochlamydia_sp.                      IS......ELPFPGLSWGLN....................................YSVS
Arabidopsis_thaliana                  VD.KHPCRHDPSHLVTHR......................................VQSN
Campylobacter_insulaenigrae           ID..KHWHPDTYFALTER......................................VKAA
Gramella_forsetii                     ID..SKKAYNTFFCLNKH......................................FETD
Burkholderia_sp.                      IA.PCRVAHDYLHQALDSRDVVVAPSASALLALGRLLGEAVRGRMPLSACAEATTEATAE
```

```
                                      β14                         α8                      α9
Mycobacterium_tuberculosis
                                      320                330       340       350       360

Mycobacterium_tuberculosis            VTT.............GAPRPAWLDRCAAMNRHAIAAVREQL.AAHPLTTGLHVAAAVSH
Mycobacterium_canettii                VTT.............GAPRPAWLDRCAAMNRHAIAAVREQL.AAHPLTTGLHVAAAVSH
Rhodococcus                           VVS.............GEPDRAWINRCRTLSEHTDKAVRAQL.AAHPKATGLHVAAAVMD
Clostridum_paraputrificum             EVS.............GECPDQWLDVCEAASELGASAVREVLGETADELTGLHVAAAVAD
Pseudomonas_aeruginosa                KTT.............GQPSKQWLKICEAASELGAEAVRDVL.AQDFSFSGLHVAAAVAD
Phaeospirillum_molischianum           PGLLADSLAAVVT...KAVPVEWLDSWYAAEQRSAVAADRL......AVPEAALFRAAIA
Sulfuricella_denitrificans            ADAVCAALVAAQP...FPAPVAWLEDFAAEEQRAAGLAQTS.....VKPIEAKVVESLIH
Afifella_pfennigii                    EAHFCRALRA......APAPAGWLKELRALDRAAGEAAEAA..CAGPQAFEGAVLRRLFQ
Tropheryma_whipplei                   EDSFSGIPLGAAK..FASASAGCIS.........QDVISSDR.PRGHRVTRQQLVSEIWN
Listeria_monocytogenes                ERFLLDIMQQNMP..DDAKDAAWLSRWTSYNKVAREIVLAEM.ANTTILEEGKIVAELRR
Bacillus_subtilis                     ASAFAEAIIMEHA..DMTKSSEWLNKWQFVNERFRAHLQTI..SSEEVSFEGNLYRILQH
Thermosynechococcus_elongatus         PQAVDCPPD.......PLPPNPYLKDWQDQDQRVREQLKRTF.EAIDWFSEVKLIYHLPQ
Clostridium_pasteurianum              PRAFAQSISI......ENSSTTYLSKWLNYQDEMRKQLNST..KEEAKLFEGKLIQSLQN
Staphylococcus_aureus                 ANDFFRSLMEDT....TVNRVNWLEKWQSLEKKGRQEIENYL...EQATDESAFVGELIK
Halobacterium_salinarum               PAATARAIAARA....DGTRSAWTDAVLDLEARYWAAVDDF...QPAATLEGEIAATVAA
Cytophaga_hutchinsonii                LNEFLSKTTIA.....SNTQEQFVQRWIQQQTHIIPAIREFN.TKEKVFNEFTAVFDVLN
Escherichia_coli                      IADWLELHP.......AEKRQPWC...VEIPRLAEQAMQAVI.ARRDAFGEAQLAHRICD
Salmonella_enterica                   IADWLELHP.......AEKRKPWC...VEIPRLAELAWQRVV.AQRDTFGEAQLAHRIRD
Haloarcula_marismortui                PDATADALAGETL...GGVAASWREQFIRAEQTHWDAVTET...ASETYWEGGVLSDVTA
Parachlamydia_acanthamoebae           LKSFFANYDV......LVSFDRAARNWIDADRQYQLAFQAL..CAEEPCAEPAMIASFSR
Natronomonas_pharaonis                ETRLATAVADAV....DRTPGSYADRLAELEPGYWRLVEGE......EPQEGAMLADAVA
Trichodesmium_erythraeum              VEQLAKILITQYF..NNNHDINYLNLWCQAEEKVRENIDTKM.AKINHILEPKISWLISQ
Syntrophus_aciditrophicus             AFAFVEGLHLQEN...PSLDATACCLFLDSLETLYQALQRR..IPEETLSFSRIASDLLG
Desulfatibacillum_alkenivorans        VGKFFTALGELKN...VSQNKEAAESLVQAAEQVRADLTAA..TPRDALSFPLIAKIINQ
Flavobacterium_psycrophilum           PSVFFKEFIPKI....NSLESNYLPYSLEIKKVRTQKTNTY..LATIPFSDFKAFEIILP
Campylobacter_blaseri                 VSCFFRQIKEF.....KKDDKSYLEIWKKTANLVEKTYQNY..LKELPFCDMSVYSEISK
Candidatus_Protochlamydia_amoebophila LREFFKNYRL......SFKPIYSFQAWKDANRSYQLELLKL..FEEEPLAELSLFHLLSK
Haemophilus_influenzae                VHHWLRAHP.......PLRQKPWLLEPLALSKFCATFIEQQV...GGNLTEASLALRLPT
Azoarcus_communis                     EAFFFERMCGFAG...AKASGTYFNAWASVERNIS........EPQATFGEIYAVGQLVR
Prochlorococcus_marinus               VDKMLEVIPVKPAIDKKIVSKKLIKELIKYDLFIHDWLDKRL.FRNGLITEPALARLLPR
Neochlamydia_sp.                      LHEFFSVYQY......SYAINKSCSSWLKEDQRYQSKLLAL..LQEEPQAEASLIYHLSK
Arabidopsis_thaliana                  IVQFANCVLKSRF...PWRRSKLHGHLQALDGAIAREMSFQI.SAESSLTEPYVAHMLSK
Campylobacter_insulaenigrae           PEVFFQQLLSVA....RLEPHSYFNLWTNLKDKADHRHTQF..LPKVPFSDFLAMSYLSK
Gramella_forsetii                     VNSFFSEFFPLT....KRTESDYGSFWKDIKGKRQHRHEDY..MAEIPYSDLKAMQEIYQ
Burkholderia_sp.                      ATAEATAHPSQPA...APVAFAWRDRWAGVASYGARERRAC..VAALEWGEVSAVHRVL.
```

```
                                   β15       α10                  β16
Mycobacterium_tuberculosis          TT                    TT          TTT
                                    370       380       390       400

Mycobacterium_tuberculosis          ALRPGDQLVLGASNPVRDVALA.GLDTRG................IRVRSNRGVAGIDGT
Mycobacterium_canettii              ALRPGDQLVLGASNPVRDVALA.GLDTHG................IRVRSNRGVAGIDGT
Rhodococcus                         ALTDGDQLLLGASNPVRDAALV.SYPAPK................IRVLSNRGVAGIDGT
Clostridum_paraputrificum           SLRTGEQLVLGASNPVRDASWA.GLPFPG................VDVHAGRGAAGIDGT
Pseudomonas_aeruginosa              TLATGDTLVLGASNPVRDASFV.GLPFSA................VDTYTPRGTAGIDGT
Phaeospirillum_molischianum         ALPEGSLLFVANSMAVRDLDDFSGSSDKR................LTVLCNRGLSGIDGN
Sulfuricella_denitrificans          QL.AGGCLFSGNSMAIRDLDNFASGGEAS................LRIIGNRGASGIDGN
Afifella_pfennigii                  ALPEETPVFLGNSLVIRAADWLAGTGPTP................LRLLGNRGLSGIDGN
Tropheryma_whipplei                 ITKEDEILFFGSSQLIRVADSI..LPGKR................VTVFANRGLSGIDGN
Listeria_monocytogenes              LLPEKAGLFIGNSMPIRDVDTYFSQIDKK................IKMLANRGANGIDGV
Bacillus_subtilis                   LVPENSSLFVGNSMPIRDVDTFFEKQERA................FRIYSNRGANGIDGV
Thermosynechococcus_elongatus       WLPSQTAIFVASSMPVRDVESVWQGSDRR................HRFYFNRGANGIDGT
Clostridium_pasteurianum            LLPSGSRLVVSNSMAIRDVDYFFEARNQN................IKVLCNRGANGIDGI
Staphylococcus_aureus               KTFEEDALFISNSMPIRDIDNL..LLNKN................IDIYANRGANGIDGI
Halobacterium_salinarum             GAPDPATVFVSNSMPIRDFDRFAAPRGAD................LAVLGNRGASGIDGV
Cytophaga_hutchinsonii              HLPAFAKLHLANSMSVRYANTL.GISQRN................AEVFANRGTSGIDGV
Escherichia_coli                    YLPEQGQLFVGNSLVVRLIDAL.SQLPAG................YPVYSNRGASGIDGL
Salmonella_enterica                 YLPEQGQLFVGNSLVVRLIDAL.SQLPAG................YPVYSNRGASGIDGL
Haloarcula_marismortui              LAPDPATLFISNSMPIRDMDRFGGPRDAD................LTVLGNRGASGIDGI
Parachlamydia_acanthamoebae         LLPRHSRMYLGNSLPIREWDMAAINEYRD................IRVEANRGVSGIEGQ
Natronomonas_pharaonis              LAPDPSTVFVSNSMPVRDLDRFGAPQAAS................LSVLGNRGASGIDGI
Trichodesmium_erythraeum            TLPKNTSIFVANSMPVRDVEFFWVPNNSQ................IQPFFNRGVNGIDGT
Syntrophus_aciditrophicus           AVPDGEGLFLGNSLVIRAFDKFRSPFPRK................ISVISNRGVSGIEGN
Desulfatibacillum_alkenivorans      NIPGGEALFPGNSNAIRAFDMAGPALAKG................VQVVTNRGVSGIEGN
Flavobacterium_psycrophilum         KLPLYSQLQISNSSAIRYAQLF..SIHSS................IEVFCNRGTSGIDGS
Campylobacter_blaseri               MLPKNSIVHLGNSSPIRYMILN..KPKDG................CKYLSNRGVSGIDGS
Candidatus_Protochlamydia_amoebophila KIPEESKIYVGNSLPIREWDQAADWTNKH................FQVFANRGANGIEGQ
Haemophilus_influenzae              LLPYNGVLFLGNSLLVRLVDAL.TQLPES................YPVYTNRGASGIDGL
Azoarcus_communis                   KLPQHAKLHIANSLPIRMVHLF..RSDPT................ITSFCNRGVNGIDGC
Prochlorococcus_marinus             LLPESIPVMIASSSPIRDWLSYSGEGAFL................RRCFGFRGASGIDGT
Neochlamydia_sp.                    QIASSSLVYLGNSLPIREWDAYATREEEH................ENIFANRGLNGIDGQ
Arabidopsis_thaliana                ALTSKSALFIGNSMPIRDVDMYGCSSENSSHVVDMMLSAELPCQWIQVTGNRGASGIDGL
Campylobacter_insulaenigrae         AIPQDYHIHFSNSSVIRYAQLF....DFK................HTIHCNRGTSGIDGS
Gramella_forsetii                   KIPKNSVLHFGNSSTIRYAQLF..EWDKS................LKIYCNRGTSGIDGS
Burkholderia_sp.                    AAPGFAFVHLGNSMSMRHADIGYDVRAAR................QDIYVNRGVSGIDGT
```

```
                                           α11                        β17  α12   η8               β18
Mycobacterium_tuberculosis                                                              TT
                                    410       420       430       440       450       460

Mycobacterium_tuberculosis          VSTAIGAALAYEGAHERTGSPDSPPRTIALIGDLTFVHDSSGLLIGPTEPIPRSLTIVVS
Mycobacterium_canettii              VSTAIGAALAYERTH...GSPDSPPRTIALIGDLTFVHDSSGLLIGPTEPIPRSLTIVVS
Rhodococcus                         VSAAVGAALAYEEG...........RTVALLGDLTFLHDASGLLIGSGEPRPRDLTIVVA
Clostridum_paraputrificum           VATAVGVALARDRAH...ADEIRPPRTLALMGDLTFLHDAGALNIGPAEPRPESLTIVVA
Pseudomonas_aeruginosa              VSQAIGVALAVQTRN...PEEIRAARTVALMGDVTFLHDVGGLLIGPDSPQPENLTIVVA
Phaeospirillum_molischianum         LSTFFGAVASGRFP...........AALALVGDLTFLHDLSGLAAGQ....GMSATICLF
Sulfuricella_denitrificans          VSTALGLCAALGKP............VVALLGDLAFYHDMNGLLAAR....GLNITFVVL
Afifella_pfennigii                  LSTAFGIAAASG.P............SLAIVGDLAFLHDLGALALGR....DLPLAILLL
Tropheryma_whipplei                 LSTAAGIAIATNRT............TRVLVGDLTGVHDIGALLIPLME.AKYRLQIVIG
Listeria_monocytogenes              VSSALGASVV.FQP............MFLLIGDLSFYHDMNGLLMAKK..YKMNLTIVIV
Bacillus_subtilis                   VSSAMGVCEGTKAP............VTLVIGDLSFYHDLNGLLAAKK..LGIPLTVILV
Thermosynechococcus_elongatus       LSTALGVAHR.GQP............TLLITGDLACLHDTNGWLITPQ..FQGCLTVLLI
Clostridium_pasteurianum            VSTALGISTS.NKP............TVLLTGDLSFYYDLNGLLIGKA..NNLNLIILLL
Staphylococcus_aureus               VSTALGMAV..HKR............VTLLIGDLSFYHDMNGLLMSKL..NNIQMNIVLL
Halobacterium_salinarum             VSSALGAGSATADP............VVGLVGDLAYFHDSNGLLALER..CGVDATIVLV
Cytophaga_hutchinsonii              ISTAYGYALKTSQL............VTIITGDLAFFYDRNAFWNNY...KPSNLRVVLL
Escherichia_coli                    LSTAAGVQRASGKP............TLAIVGDLSALYDLNALALLRQ..VSAPLVLIVV
Salmonella_enterica                 LSTAAGVQRASAKS............TLAIVGDLSALYDLNALALLRQ..VSAPFVLIVV
Haloarcula_marismortui              TSTALGAGSATSDP............LVLVTGDLAYYHDMNGLLALGR..CAVDATVVLL
Parachlamydia_acanthamoebae         ISTFLGWCIPNTP.............QFALLGDLTTLYDLAGPWILHQ.LQNVSAQIIVI
Natronomonas_pharaonis              ASTALGAGFGTDDP............LVAVTGDLAYYHDMNGLLAVSR..AGVDATIVCI
Trichodesmium_erythraeum            LSTALGIAHRYQK.............TVMLTGDLALLHDTNGFLLRNK..LVGHLTIILI
Syntrophus_aciditrophicus           IATSVGFAEASRRR............VTAVIGDISFLHDLNSLLLLAQ..SATPVVLIII
Desulfatibacillum_alkenivorans      LATSLGFAEGAGKR............VTAVIGDVSMLHDLNSLAMVSA..SRAQVITIIV
Flavobacterium_psycrophilum         TSTAIGAAIASGKE............TILITGDISFLYDSNALWNNY...IPENFKIILL
Campylobacter_blaseri               FSSAVGYASKDEKI............NTVILGDLSFLYDSNAMWNGY...LKENLKIILV
Candidatus_Protochlamydia_amoebophila LSTFLGLASPEQN.............NWAILGDLTTLYDMIAPWVIHQ.IQPLKVNVVVV
Haemophilus_influenzae              LATAAGIGIGSNKP............VVAVIGDTSTLYDLNSFALFKN..VTQPTLIFVI
Azoarcus_communis                   MSTAVGYAAASAEP............VYLVIGDLTFFYDMNALWNRH...LSKSLRILLL
Prochlorococcus_marinus             LSMGMGLSIIMGR.............MVLVTGDLALLHDTNGWLFSKD..KNISLIVIMI
Neochlamydia_sp.                    LSTFLGMCSCRQQ.............NWGIMGDLTTLYDLAAPWILEQ.LKEIDLNLLVI
Arabidopsis_thaliana                LSSATGFAVGCKKR............VVCVVGDISFLHDTNGLAILKQRIARKPMTILVI
Campylobacter_insulaenigrae         TSTAMGYAMVSDSP............TVLITGDIGFFYDINGLWNQY...IPPYTRIILL
Gramella_forsetii                   VSTAVGASVSSEEP............VTIITGDLSFFYDSNALWNNY...IPSNFRIIIL
Burkholderia_sp.                    LSTFIGEALARRDP............GLLLLGDQALVHDLSALASAQR..VRTPACVCVV
```

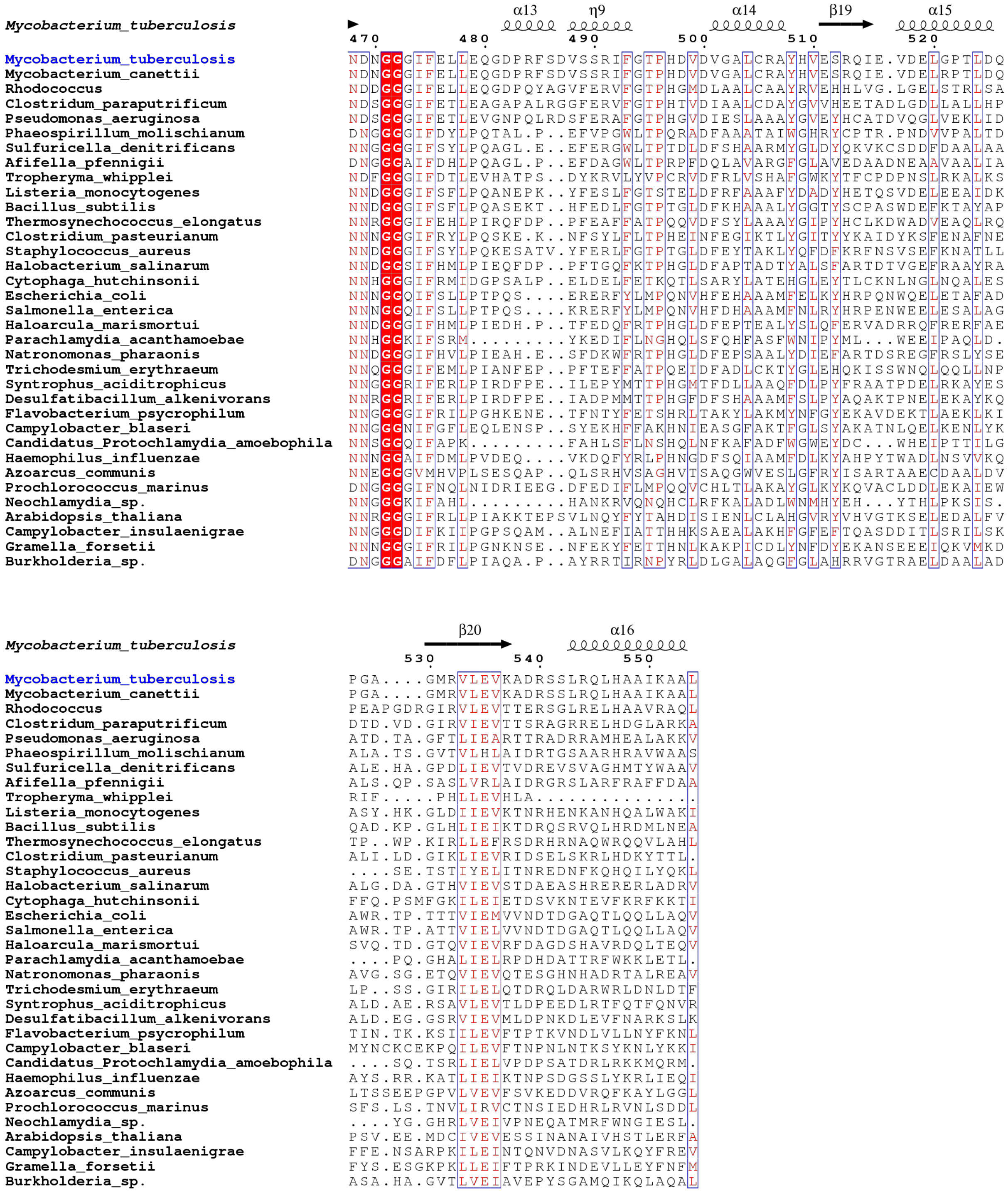


**Supplementary Figure 1 (SF1) Caption**: MAFFT alignment of 35 bacterial, archaeal and plant MenD sequences. Figure generated using ESPript (reference in main text). Conserved residues are shown in red boxes (strictly conserved) or red text. Arginine residues at the 97, 277, 303 and 305 ("WR") position are highlighted using yellow boxes, with arginine-like residues (His, Lys) at these positions shown as blue text.
